# Transcriptional Regulation of Microglial Metabolic and Activation States by P2RY12

**DOI:** 10.1101/2025.07.21.665960

**Authors:** Aida Oryza Lopez-Ortiz, Madison Doceti, JaQuinta Thomas, Abigayle Duffy, Morgan Coburn, Akhabue Okojie, Audrey Lee, Elizabeth Aidita Sou, Alban Gaultier, Ukpong B. Eyo

## Abstract

Microglia are the resident immune cells of the CNS. Under homeostatic conditions, Microglia play critical roles orchestrating synaptic pruning, debris clearance and dead cells removals. In disease, they are powerful mediators of neuroinflammation, as they rapidly respond to injury or infection within the CNS by altering their morphology, proliferating, and releasing cytokines and other signaling molecules. Understanding the molecular pathways involved in microglial function is pivotal for advancing neurobiological research and developing effective strategies for CNS disorders. In this context, P2RY12 is a G protein-coupled receptor (GPCR) that is uniquely enriched in microglia in the parenchyma and a canonical marker of homeostatic, ramified microglia. However, P2RY12 is downregulated in activated microglia and in neurological conditions. The consequences of P2RY12 downregulation in disease-associated microglia and how they influence microglial activation remain poorly understood. In this study, we apply transcriptional and histological methods to explore the changes to microglia upon a genetic P2RY12 loss. Our findings reveal that P2RY12-deficient microglia experience alterations in distinct metabolic pathways while preserving overall homeostatic microglial transcriptional identity. Lack of P2RY12 alters signature genes involved in homeostatic iron metabolism. Importantly, the genes encoding proteins in the Glutathione Peroxidase 4 (*Gpx4*)-Glutathione (GSH) antioxidant pathway related to ferroptosis susceptibility are impaired upon microglial activation with lipopolysaccharide (LPS) treatment. These results highlight the critical role of P2RY12 in regulating microglial immune and metabolic transcriptional responses under both homeostatic and inflammatory conditions, providing insights into its involvement in CNS pathophysiology.

**Proposed Model:** 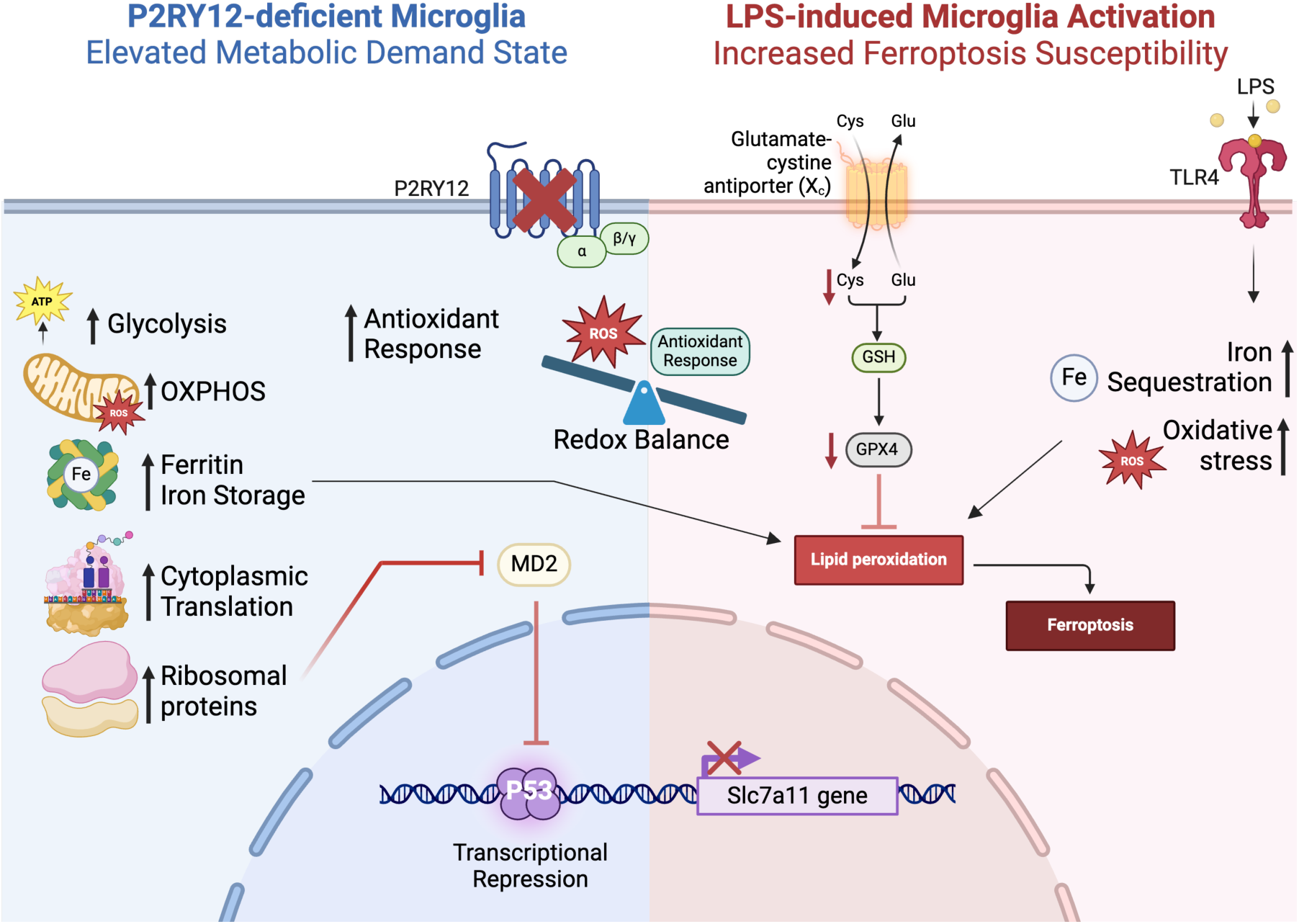

At basal condition the transcriptional landscape of P2RY12-deficient microglia suggests that the cell is in a highly demanding metabolic state with increased oxidative phosphorylation and glycolysis as well as augmented expression of ribosomal proteins involved in cytoplasmic translation. Additionally, P2RY12-deficient microglia exhibit heightened expression of key regulators of the antioxidant response, suggesting elevated ROS levels as a consequence of this highly metabolically active and energetically demanding microglial state. Elevated ROS levels might diminish microglial antioxidant reserves thus rendering them more vulnerable to additional oxidative stress pushing the redox balance. When activated by LPS treatment, P2RY12-deficient microglia exhibit an impaired responsiveness of system x_c_ and downstream enzymes involved in GSH mediated antioxidant response that together with LPS induced intracellular iron sequestration(François et al., 2014; S. Liu, Gao, & Zhou, 2022; McCarthy et al., 2018; Sfera et al., 2018; Shin et al., 2018; Urrutia et al., 2013) suggesting that P2RY12-deficient microglia will show increased susceptibility to ferroptosis. It is possible that system x_c_ unresponsiveness in KO microglia results from the upregulation of ribosomal proteins involved in the inhibition of MD2, a negative regulator of p53 DNA binding activity(Daftuar, Zhu, Jacq, & Prives, 2013). In this sense, the repression of p53 inhibition will enhance its function, possibly explaining a suppression of *Slc7a11* expression in LPS-activated P2RY12 deficient microglia.

## Introduction

Microglia are the resident innate immune cells of the central nervous system (CNS). They play pivotal roles in tissue homeostasis as well as degeneration(Salter & Stevens, 2017; Sierra, Miron, Paolicelli, & Ransohoff, 2024) including immune functions that contribute to normal brain function, supporting neurogenesis(Diaz-Aparicio et al., 2020; Sierra et al., 2010), synapse monitoring (Paolicelli et al., 2011; Tremblay et al., 2011), glutamatergic synapses regulation(Basilico et al., 2022), neuronal excitability(Badimon et al., 2020; Basilico et al., 2022; J. Peng et al., 2019; Umpierre & Wu, 2021), memory (C. Wang et al., 2020) as well as cerebrovascular regulation(Bisht et al., 2021; Császár et al., 2022; Fu et al., 2025). The fine tuning of microglial activation relies heavily on immunometabolic reprogramming(Bernier, York, & MacVicar, 2020), where it has been documented that upon activation under normoxia, microglia undergo a metabolic shift from oxidative phosphorylation to glycolysis, known as the “Warburg effect”(Sabogal-Guáqueta et al., 2023). Besides a more rapid production of energy in form of ATP, this shift allows the generation of intermediaries for cellular biosynthesis of nucleotides, amino acids and lipids required to support cellular processes associated with microglial activation such as phagocytosis, cell motility, and proliferation, as well as increased microglial production of cytokines, nitric oxide (NO), and reactive oxygen species (ROS) needed for pathogen defense(Hanisch & Kettenmann, 2007; Kettenmann, Hanisch, Noda, & Verkhratsky, 2011; Laroux, Romero, Wetzler, Engel, & Terhorst, 2005; Perry & Holmes, 2014; Perry, Nicoll, & Holmes, 2010; Ransohoff & Perry, 2009). However, disruptions in this metabolic transition can result in prolonged activation, leading to mitochondrial dysfunction and exacerbated oxidative stress that alters inflammatory responses(Baik et al., 2023; Houldsworth, 2024). This can result in dysregulated neuroinflammation, influencing the progression of neurological and neurodegenerative disorders(Socodato & Relvas, 2025). Given their essential functions, understanding the molecular mechanisms underlying microglial homeostasis and activation is crucial.

Microglia have a distinct ontogeny from other CNS cells where they enter the brain embryonically in mice and develop in the CNS unique environment, protected from the periphery by the brain blood barrier(Ginhoux et al., 2010; Ginhoux & Prinz, 2015; Obermeier, Daneman, & Ransohoff, 2013). Microglia presents with a distinct gene expression profile that distinguish them from other peripheral immune cells as well as from other brain cells(Bennett et al., 2018; Cronk et al., 2018; Lavin et al., 2014). Amongst microglial signature genes, the *P2ry12* gene encodes the P2RY12 G protein-coupled receptor, which is part of the microglial sensome(Hickman et al., 2013) enabling microglia to respond to ATP secreted in both physiological and pathological conditions(Gómez Morillas, Besson, & Lerouet, 2021)*. P2ry12* is primarily expressed in ramified, surveillant microglia in the healthy brain(Haynes et al., 2006; Mildner, Huang, Radke, Stenzel, & Priller, 2017; Moore et al., 2015). However, under pathological conditions, *p2ry12* / P2RY12 is significantly downregulated and is linked with activated states, including when isolated in culture or in disease-associated microglia (DAM) in pathology (Bohlen et al., 2017; H Keren-Shaul et al., 2017). These activated states have been described in response to neurodegenerative signals like amyloid plaques, apoptotic bodies, or myelin debris(A. Deczkowska et al., 2018). Similarly, the DAM phenotype is present in response to PAMPs such as bacterial lipopolysaccharide (LPS), a ligand for Toll-like receptors(Lively & Schlichter, 2018; Rangaraju et al., 2018). While *P2ry12* / P2RY12 downregulation is used as an indicator of a loss of homeostatic microglial states(van Wageningen et al., 2019; Zrzavy et al., 2017), the intracellular consequences of its downregulation and how it influences microglial transformation into activated phenotypes, particularly under inflammatory conditions remains unknown.

In the present study, to elucidate the role of *P2ry12* in the regulation of the microglial homeostatic state and responses or inflammatory stimuli, we evaluated the transcriptional profile in microglia following genetic deletion of *P2ry12* / P2RY12. We found that while maintaining a general microglial identity, a *P2ry12* / P2RY12 deficiency shifted microglial transcription toward an altered immunometabolic and hypermetabolic state. Interestingly, upon LPS challenge *P2ry12* / P2RY12 deficient microglia exhibited alterations in genes that code for proteins associated with the GPX4-Glutathione (GSH) antioxidant response consistent with glutathione depletion and increased ferroptosis susceptibility. Our findings provide new insights into possible *P2ry12* / P2RY12 regulation of microglial immunometabolic programming and activation. A better understanding of P2RY12’s role in microglial immunometabolic regulation may clarify the link between neuroinflammation, oxidative stress, and CNS disorders.

## Methods

### Animals

Both male and female mice in this study were on a C57BL/6J genetic background. Littermate mice were generated by pairing male and female P2RY12 heterozygous mice generated from crossings with wildtype (WT) and P2RY12KO (KO) mice. All animal experiments were approved and complied with regulations of the Institutional Animal Care and Use Committee at the University of Virginia (Protocol No. 4237).

### LPS challenge

Mice were given a single intraperitoneal LPS injection at a 1 mg/kg dose to induce mild neuroinflammation. Changes in the internal temperature and weight were measured 6 and 24 hours after the treatment and the end point was reached at 24h.

### Microglia Isolation

Mice were euthanized with CO_2_ and transcardially perfused with 30mL ice cold 1X phosphate buffered saline (PBS) + Heparin 0.05% (wt./vol, Product #: 00409-2720-02; Pfizer Inc, USA). The brain was immediately dissected from cerebellum and grossly minced to pieces with a scalpel and suspended in 3mL of HBSSinh: Hanks’ Balanced Salt Solution (HBSS) without Ca^2+^ and Mg^2+^ (Product #: 14175095; Thermo Fisher Scientific, USA) that included an inhibitor cocktail to prevent microglial activation: we used the transcription inhibitors Actinomycin D (5µg/mg, w/w, Product #: A1410; Millipore Sigma, USA) and Triptolide (10µM, mol/L) (Product #: T3652; Millipore Sigma, USA) in addition to the translational inhibitor Anisomycin (27.1µg/mL, w/w) (Product #: A9789; Millipore Sigma, USA) adapted from(Marsh et al., 2022). The brains were placed in ice until all the brains were collected. Afterwards, we proceeded with enzymatic digestion of the brains by adding papain (4U/mL) (Product #: LS003126; Worthington Biochemical Corp, USA) and DNaseI (50U/ml) (Product #: E1011; Zymo Research, USA) to each sample just before starting enzymatic digestion. Samples were placed in a warm bath and incubated for 15 min at 37°C, then triturated using an 18-gauge needle, repeating 3 times without exceeding 45 minutes. Using the same syringe, the samples were passed through a pre-soaked 70µm filter into a 50mL conical tube washing afterwards with 5mL HBSSinh and spun at 1500rpm at 4°C for 10 minutes. For myelin removal, the supernatant was vacuumed, and the resulting pellet was resuspended in 20mL of Percoll 40% (wt./vol, Product #: 17-0891-02 ; Danaher Corporation, USA) + the inhibitor cocktail. The percoll suspension was centrifuged at 450xg for 25 min at 4°C with no break. The myelin-containing supernatant was completely removed, and the cellular pellet was resuspended in 180µl of FACS buffer (1X PBS + Bovine Serum Albumin 0.2% (wt./vol, Product #: 130-091-376; Miltenyi Biotec Co, USA) + EDTA 0.5µM (mol/L, Product #: AM9912; Thermo Fisher Scientific, USA) containing Actinomicyn D. To separate CD11b-positive microglial cells, 20µL of anti-CD11b-Biotin microbeads (Product #:130-049-601; Miltenyi Biotec Co, USA) were added to the resuspended pellet and incubated for 15 minutes at 4°C degrees followed by magnetic separation with LS columns (Product #:130-042-401; Miltenyi Biotec Co, USA) following the manufacturer’s instructions.

### RNA extraction

RNA was isolated from CD11b-positive microglial cells by using Direct-zol RNA Miniprep Plus (Product #: R2072; Zymo Research, USA) according to manufacturer’s instruction. The isolated RNA was immediately frozen at -80°C and sent to sequence at Yale Center for Genome Analysis (YCGA, New Haven, Connecticut) and UVA Genome Analysis and Technology Core. The RNA quality in the samples was measured using the Agilent 2100 Bioanalyzer with the Eukaryote Total RNA Pico Assay. The RNA Integrity Number (RIN) was used as a quality metric, and samples with values greater than 7 were sequenced (**Supplemental Table 8**).

### Bulk RNA Sequencing

Two bulk RNA sequencing datasets were generated using different protocols. For the first dataset (Basal), libraries were prepared using the NEBNext® Ultra™ II Directional RNA Library Prep Kit for Illumina (Product #: E7760L, New England Biolabs, USA) following poly(A) mRNA enrichment using the NEBNext Poly(A) mRNA Magnetic Isolation Module (Product #: E7760L, New England Biolabs, USA). Sequencing was performed on an Illumina NextSeq 2000 system. For the second dataset (LPS), RNA libraries were prepared using the KAPA RNA HyperPrep Kit (Product #: KR1352, Roche Molecular Systems, USA) with poly(A) selection, and samples were sequenced on an Illumina NovaSeq X platform. Raw sequencing data from both datasets were used for downstream transcriptomic analyses. Raw sequencing data from both datasets were deposited in the NCBI Gene Expression Omnibus (GEO) under accession number GSE296508 and used for downstream transcriptomic analyses.

### RNA Sequencing Data Processing

Sequencing quality was assessed using Fastqc (https://www.bioinformatics.babraham.ac.uk/projects/fastqc/). Next, the bioinformatic tool cutadap (Martin, 2011) was used to remove adapter sequences, trim low-quality bases, and filter short reads. Lastly, processed reads were mapped to the mouse reference genome (GRCm39) using STAR v2.6.1(Dobin et al., 2013) and reads that map onto each gene will be quantified using the featureCounts tool included in Subread software package (https://subread.sourceforge.net/). The generated count table was imported to R to perform differential gene expression (DEG) analysis using the DESeq2 package(Love, Huber, & Anders, 2014). Lowly expressed genes were filtered out, retaining genes with at least 15 counts in 14 or more samples to improve data robustness. Variance stabilizing transformation (VST) was applied to normalize the dataset for downstream analysis.

### Weighted Gene Co-Expression Network Analysis (WGCNA)

To identify gene co-expression networks, WGCNA was performed on the variance-stabilized expression data using the R WGCNA package(Langfelder & Horvath, 2008). Prior to network construction, excessive missing values and outlier samples were assessed using the function goodSamplesGenes. Soft-thresholding power selection was determined using the function pickSoftThreshold, and a signed network was constructed using the function blockwiseModules with a soft power of 12. Module eigengenes were correlated with experimental traits (genotype and sex) using Pearson correlation, and statistical significance was assessed via corPvalueStudent. Modules significantly associated with P2RY12-deficient microglia were selected for further functional analysis.

### Cytoscape Network Visualization

The top co-expressed genes from the cyan and pink modules were extracted, and adjacency matrices were calculated using TOM similarity to determine interaction strength. Highly connected genes (threshold ≥ 0.8) were exported to Cytoscape(Shannon et al., 2003) for network visualization, enabling the identification of potential hub genes within microglial P2RY12-dependent pathways. Additionally, the top 50 nodes ranked by Maximal Clique Centrality (MCC) were identified. MCC is a network centrality measure that is particularly useful in detecting key regulatory nodes or hub genes essential for maintaining the structure and function of biological networks, such as gene regulatory networks.

### Functional Enrichment Analysis

Gene Ontology (GO) was performed using <u>ShinyGO 0.82</u>(Ge, Jung, & Yao, 2020) selecting the Biological Pathway and Molecular Function databases. Additionally, GO pathway analysis was performed using the function enrichGO from the ClusterProfiler R package(G. Yu, Wang, Han, & He, 2012) to identify enriched biological processes (BP) for the WGCNA. Bar plots of enriched GO terms were generated to visualize overrepresented pathways.

### Immunofluorescence

Mice were first perfused with 1X PBS (Life Technologies Corporation, NY, USA) + Heparin 0.05% and then by 4% (wt./vol) paraformaldehyde (PFA, Product #: 158127; Sigma-Aldrich, Germany) in 1X PBS, after which they were decapitated, and their brains harvested and post-fixed for less than 48h in 4% (wt./vol) PFA. Then the brains were washed in PBS and stored in a 30% sucrose solution at 4°C. Brains were cryopreserved in optimal cutting temperature compound (OCT) media (Product #: 72592; Electron Microscopy Science, USA) and then sectioned in 30µm thick free-floating sections were obtained using a cryostat (Model CM1950, Leica). Four sections per mouse were incubated in blocking buffer consisting of 1% Bovine Serum Albumin (BSA, Sigma-Aldrich, Germany), 10% normal donkey serum (Product #: 017-000-121; Jackson ImmunoResearch Laboratories, Inc. USA) and 0.3% triton X-100 solution (Product #: 93443; Sigma-Aldrich, Germany) in 1X PBS for 1h at room temperature. Anti-PGK1 (Rabbit anti-PGK1, Product #: PA5-28612, Thermo Fisher Scientific, USA) and Anti-IBA1(Goat anti-IBA1, Product #: 011-27991, FUJIFILM Wako Pure Chemical Corporation, Japan) primary antibodies were prepared in blocking buffer at 1:500 dilution for both PGK1 and IBA1 antibody, and sections were incubated with primary antibodies overnight at 4°C. Secondary antibodies, donkey anti-rabbbit Alexa Fluor 405 and donkey anti-rabbit Alexa Fluor 594 (Product #: ; Jackson ImmunoResearch Laboratories, Inc. USA) were also prepared in blocking buffer at a 2.7µL per ml dilution. Sections were then incubated in secondary antibody for 2h at room temperature. Cell nuclei were stained by incubating sections in NucGreen (1 drop per mL of PBS 1X, Product #: R37109, Thermo Fisher Scientific, USA) for 5 min. Images were acquired using STELLARIS 5 confocal microscope (Leica Microsystems, USA) with a 63× objective. PGK1 area inside microglia was quantified using Fiji (ImageJ) software.

### Statistical analysis

All data are expressed as mean ± SEM. Statistical analyses were conducted using GraphPad Prism (v9.5.1 for Mac, GraphPad Software, La Jolla, CA, USA) and R (v4.4.2). Comparisons between two independent groups were performed using a two-tailed Student’s t-test, while analyses involving more than two groups were conducted using two-way ANOVA. Where applicable, post-hoc analyses were performed using Tukey’s or Sidak’s multiple comparisons tests. A p-value < 0.05 was considered statistically significant.

## Results

### *P2ry12* DEFICIENCY DOES NOT INDUCE DISEASE-ASSOCIATED MICROGLIAL TRANSCRIPTIONAL STATE

To understand the impact of a *P2ry12* gene deficiency on the microglial transcriptional landscape, we performed bulk RNA sequencing (RNAseq) on isolated microglia from adult male and female WT and KO littermate mice (**Fig 1a**). We compared male and female microglial transcriptional profiles by performing Principal Component Analysis (PCA) where microglial samples clustered according to sex (**Supplemental Fig 1b**). By assessing the differentially expressed genes (DEGs) between male and female microglia from both WT and KO, we found significantly altered genes between male and female microglia. These genes are sex chromosome linked genes such as the female-specific *Xist*(Villa et al., 2018) and male-specific *Uty*, *Kdm5d*, *Ddx3y*, and *Eif2s3* consistent with previous work(Guneykaya et al., 2018; M. Zhang et al., 2021) (**Supplemental Fig 1c**). Compared to other brain cells, microglia possess the most sexually dimorphic gene numbers(M. Zhang et al., 2021), which explains why sex is a strong clustering variable in microglia even in the absence of P2RY12. Furthermore, we evaluated the interaction effect between sex and genotype thus testing if the effect of genotype (KO vs. WT) on gene expression differences between males and females is significant (**Supplemental Fig 1c**). The 2 labeled genes (*Ighv1-18* and *Igkv19-93*) show significant (FDR < 0.05) and are highly different between males and females. Their positions on the left suggest that the KO effect is stronger in females than in males for these genes. The rest of the genes (gray dots, 291 DEGs) showed non-significant difference, meaning their KO effect is similar between sexes. As previously described, these results suggest that microglia display strong sex-specific transcriptional profiles, with samples clustering by sex and showing differential expression of sex chromosome-linked genes. Sex remains a dominant factor in microglial gene expression upon *P2ry12* deletion, moreover interaction analysis implies that the effect of a *P2ry12* loss is consistent between sexes for most genes, with only two genes (*Ighv1-18* and *Igkv19-93*) showing a significantly differential expression in females, indicating limited sex-specific genotype effects.

**Figure 1.**
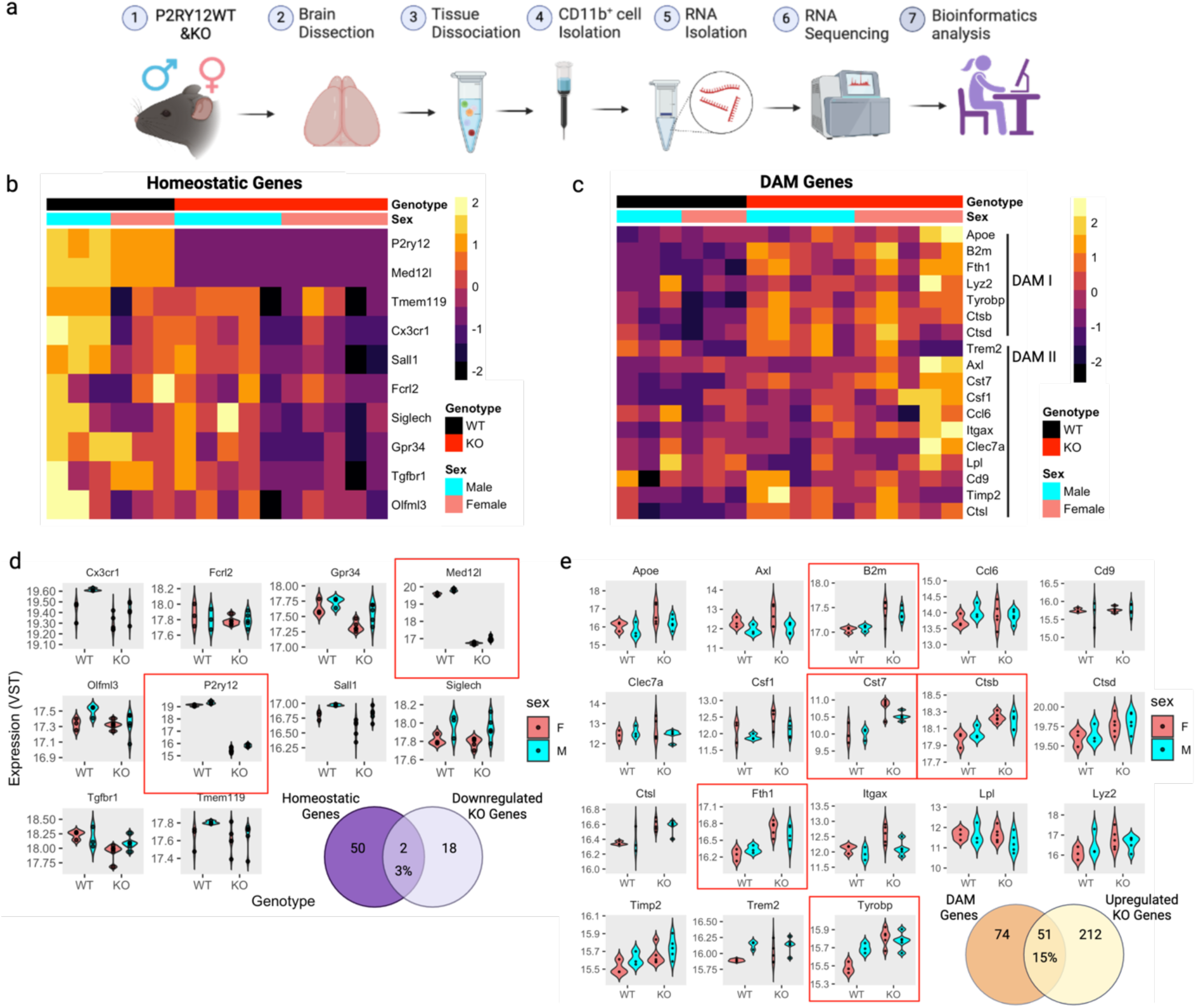
A *P2ry12* deficiency has limited effects on homeostatic and disease-associated microglial transcriptional phenotypes independent of sex. **(a)** Schematic representation of the study design. Microglia were isolated from male and female WT and KO littermate mouse brains followed by tissue dissociation, CD11b^+^ cell isolation, RNA extraction, sequencing, and bioinformatics analyses. **(b)** Heatmap displaying the expression of homeostatic microglial genes in WT and KO littermate mice, further stratified by sex. **(c)** Heatmap depicting expression levels of upregulated DAM genes in WT and KO littermate microglia as categorized into Stage 1 DAM and Stage 2 DAM subgroups by (H Keren-Shaul et al., 2017). **(d)** Distribution of homeostatic genes in WT and KO littermate microglia, separated by sex. The embedded Venn diagram compares the number of homeostatic genes reported in (Cronk et al., 2018) with significantly downregulated genes in KO mice. **(e)** Distributions of upregulated DAM-associated genes in WT and KO littermate microglia. The embedded Venn diagram compares upregulated DAM genes with significantly upregulated KO genes. Data presented as variance-stabilized transformed (VST) counts with visualizations such as heatmaps and violin plots to accurately depict expression distributions. n = 6 WT (3 male and 3 female) and 10 KO (5 male and 5 female) mice. The following tests were performed for statistical analysis: Negative binomial GLM with the Wald test for differential expression, Benjamini-Hochberg correction to control false discoveries, apeglm shrinkage to reduce variability in LFC estimates and improve biological interpretability.

Next, we explored the expression of microglial homeostatic and disease-associated (DAM) signature genes upon loss of *P2ry12* (**Fig 1b-c**). We found that amongst the previously described 52 homeostatic genes from the microglial homeostatic gene list(Cronk et al., 2018), *P2ry12* and *Med12l* were the only genes to be significantly downregulated in the P2RY12-deficient microglia when compared to their WT littermates. This represents an ∼ 96% overlap with the microglial homeostatic genes (**Fig 1d**).

Next, we compared shared genes between upregulated DEGs in P2RY12-deficient microglia and upregulated genes in the 125 upregulated DAM signature gene list(H Keren-Shaul et al., 2017). Our analysis reveals limited overlap between our dataset and the previously described DAM genes, with 51 genes in common representing about a 15% overlap (**Fig 1e**). Furthermore, we found 4 upregulated DEGs associated with the DAM1 stage: *B2m*, a component of the major histocompatibility complex (MHC) class I described in neuropathology and during brain aging(Aleksandra Deczkowska et al., 2017; Smith et al., 2015), the cathepsin *Ctsb* related to lysosomal and proteolytic activity (McGlinchey & Lee, 2015; J. Wang et al., 2021), the ferritin heavy chain encoding gene *Fth1* involved in cellular iron storage and metabolism(N. Zhang, Yu, Xie, & Xu, 2021) and lastly *Tyrobp* which encodes for a transmembrane immune signaling adaptor important for microglia’s intracellular signaling(Haure-Mirande, Audrain, Ehrlich, & Gandy, 2022). In contrast, only one gene, *Cst7*, was shared with the DAM2 stage, which is involved in the inhibition of cysteine proteases (Hamilton, Colbert, Schuettelkopf, & Watts, 2008).

Overall, these results suggest that despite a deficiency of the microglial homeostatic *P2ry12* gene, KO microglia hold a transcriptional profile compatible with homeostatic microglia at ∼96% overlap with only about ∼15% of the upregulated DAM signature overlap, suggesting that *P2ry12* deficient microglia is in a largely homeostatic, though perhaps mildly activated, state. Because our initial analyses did not find important sex differences with a *P2ry12* deficiency, we conducted the rest of our assessments in the next studies using pooled male and female data.

### DIFFERENTIALLY EXPRESSED GENES (DEG) AND PATHWAY ANALYSES IN WT AND KO MICROGLIA REFLECT IMMUNO-METABOLIC DISTURBANCES

To gain a deeper understanding of the effect of a *P2ry12* deficiency, we performed differentially expressed gene (DEG) analysis. The DEGs between the KO and WT are represented in a volcano plot (**Fig 2a**). Applying a log fold change cutoff of -1 and 1, with an FDR below 0.05, we found 283 DEGs (20 downregulated and 263 upregulated genes) in the KO microglia when compared to the WT counterparts. To better understand the biological significance of the DEGs that we found in the KO microglia, we took the list of significantly downregulated genes (without including *p2ry12*) and performed gene ontology (GO) molecular function analysis. We found that the most enriched terms relate to peptidase activity which groups the *Adamts13, Adamts16 and adamtsl2* genes (**Supplemental Table 1**). *Adamts13* and *Adamts16* encode members of the ADAMTS family of metalloproteinases, while *Adamtsl2* encodes a member of the ADAMTS-like proteins. These enzymes are involved in extracellular matrix (ECM) turnover and remodeling (Gao et al., 2012; Stanton, Melrose, Little, & Fosang, 2011). Other enriched processes include G protein-coupled receptor and purinergic activity, suggesting potential disruptions in GPCR signaling pathways (**Fig 2b**). These results suggest that downregulated genes are primarily involved in extracellular matrix remodeling, and proteolytic processes, which may have implications for microglial intracellular communication and structural integrity indicating that P2RY12 homeostatically promotes this functional state in microglia.

**Figure 2.**
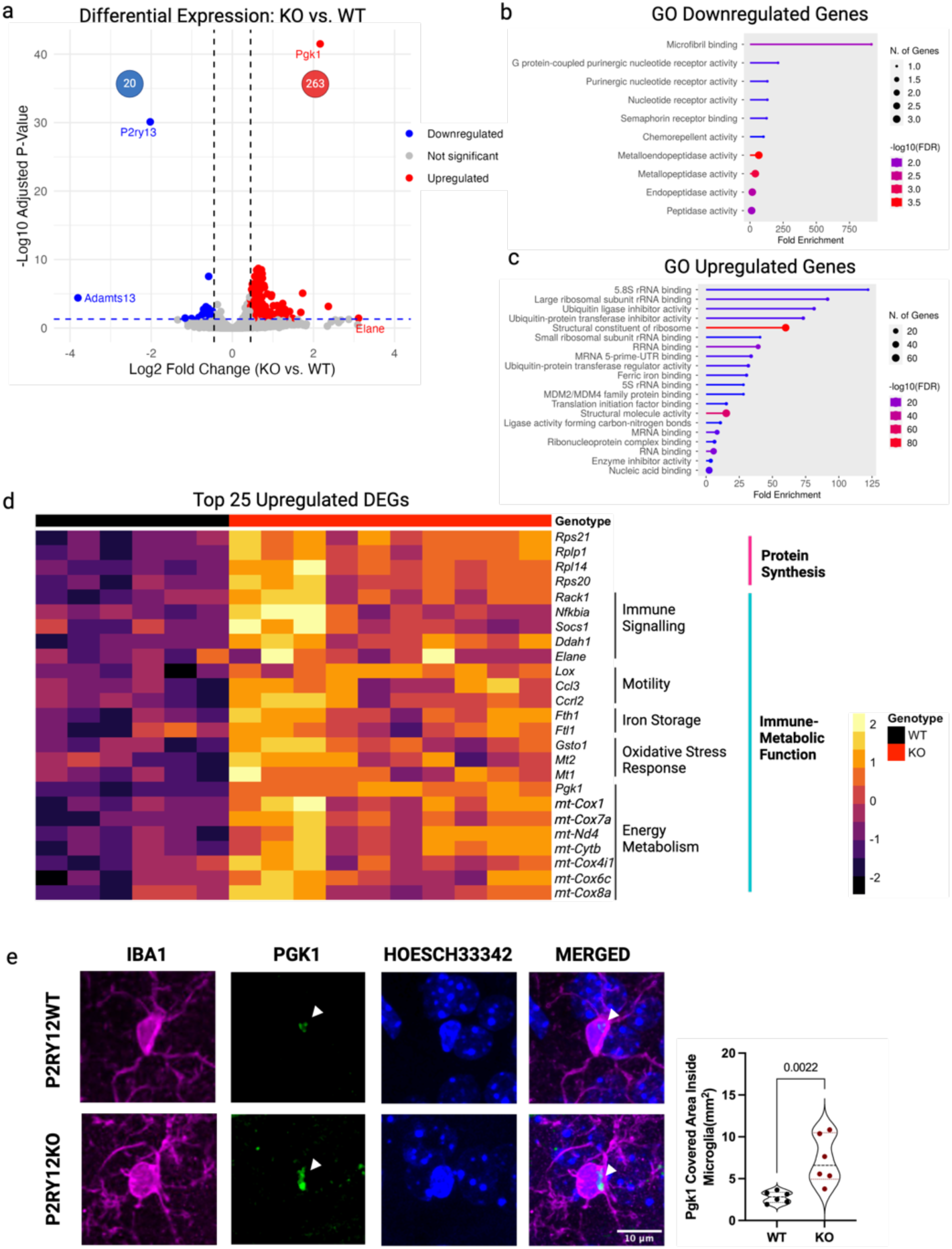
DEGs and pathway analyses in WT and KO microglia reflect immuno-metabolic disturbances. **(a)** A volcano plot showing differentially expressed genes (DEGs) in KO microglia compared to WT littermate microglia. Upregulated genes (red) and downregulated genes (blue) are indicated. (**b-c**) Gene Ontology (GO) molecular function analysis of **(b)** downregulated and **(c)** upregulated genes in P2RY12KO microglia. Functions related to GPCR signaling and extracellular matrix organization are enriched in downregulated genes, while translational regulation and iron storage are enriched in upregulated genes. **(d)** Heatmap of significantly upregulated genes in P2RY12KO microglia. Genes are categorized into protein synthesis & ribosomal function (magenta) and immune-metabolic function (cyan). **(e)** Immunofluorescence staining (left panel) of microglia (IBA1, magenta) and PGK1 (green) in WT and P2RY12KO littermate microglia. The merged panel includes DAPI (blue) for nuclear staining. The quantification of PGK1-covered area per microglial cell (middle panel) shows a significant increase in KO microglia (P = 0.0022). Data presented as mean ± s.e.m. n = 6 WT (3 male and 3 female) and 6 KO (3 male and 3 female) mice for (e). Mann-Whitney U test was used for statistical analysis. Bulk RNAseq data presented as variance-stabilized transformed (VST) counts with visualizations such as PCA plots, heatmaps and violin plots to accurately depict expression distributions. n = 6 WT (3 male and 3 female) and 10 KO (5 male and 5 female) mice. The following tests were performed for statistical analysis: Negative binomial GLM with the Wald test for differential expression, Benjamini-Hochberg correction to control false discoveries, apeglm shrinkage to reduce variability in LFC estimates and improve biological interpretability.

We repeated this process with the list of significantly upregulated genes in the KO microglia and found the most significantly enriched molecular functions to be ribosomal and translational activity (**Fig 2c**). Other functions include ferric iron binding and genes encoding MDM2/MDM4 family protein binding (**Supplemental Table 2**), associated with P53 post-translational regulation(Daftuar et al., 2013). These findings indicate that upregulated genes are primarily involved in translational processes, iron metabolism and p53 signaling in *P2ry12* deficient microglia implying that in the homeostatic state, P2RY12 may function to dampen these processes in microglia.

Since the upregulated genes were more abundant in KO microglia, we visualized the top 25 most upregulated genes in a heatmap (**Fig. 2d**). We observed that they clustered into two main functional groups. The first is the *protein synthesis and ribosomal function* group. The ribosomal function category includes genes involved in ribosomal activity and translation. The second is the *immune-metabolic function* group. This group of genes can be further subclassified in 5 categories according to their associated functions: (1) immune signaling include genes such as *Nfkbia*, a member of the NF-κB inhibitor family(T. Zhang et al., 2022), described to have a role in microglia activation(Lei & Liu, 2024; Ye et al., 2025), and *Ddah1*, whose function is associated with increasing nitric oxide (NO) production, an important mediator of macrophages / microglia signaling and function(Kopec & Carroll, 2000; Monsonego, Imitola, Zota, Oida, & Weiner, 2003; Nakanishi et al., 2001); (2) the motility associated group of genes comprises *Lox* and *Ccrl2* implicated in leukocyte migration, including in macrophages / microglia (Cordoba et al., 2021; Yang Liu et al., 2024; Schioppa et al., 2020); (3) we found the upregulation of *Fth1* and *Ftl* that encode for ferritin heavy and light chain respectively, components of the major cellular iron storage(Bou-Abdallah, Paliakkara, Melman, & Melman, 2018); (4) we also found genes such as *Gsto1*, Mt1 and Mt2 involved in oxidative stress response through cellular detoxification(Allen et al., 2012; Board et al., 2001; Couto, Wood, & Barber, 2016) and metal ion buffering mechanisms likely aimed at mitigating redox imbalance; (5) the last category includes genes involved in energy metabolism and ATP production such as the mitochondrial genes *mt-Cox1* and *mt*-*Nd4*, as well as the nuclear-encoded gene *mt*-*Cox7a2l* components of the mitochondrial respiratory chain, involved oxidative phosphorylation (OXPHOS)(L.-Y. Peng et al., 2014).

Additionally, the *Pgk1* transcript was among these metabolic related DEGs **(Fig 2a, d)**. *Pgk1* encodes the phosphoglycerate kinase 1 (PGK1) protein, the first ATP-generating enzyme in the glycolysis pathway(Nie et al., 2020). Notably, PGK1 had been recently described to be increased and associated with a pro-inflammatory microglial profile and increased glycolysis in MCAO rat models(Cao et al., 2023). To validate this finding at the protein level, we conducted immunohistochemistry for PGK1 on tissues from WT and KO littermate brains. Here, we found PGK1 is increased at the protein level within P2RY12 deficient microglial cells compared to WT microglia from littermate mice (**Fig 2e**), thus corroborating some of our transcriptional findings at the protein level.

Together, these findings provide a comprehensive transcriptional landscape of *P2ry12*-deficient microglia, revealing significant alterations in general metabolism, iron metabolism, immune signaling, and cellular homeostasis. Enriching functions related to protein synthesis, iron storage, and energy metabolism suggests that a *P2ry12* deficiency drives a distinct microglial expression profile characterized by enhanced protein synthesis and energy production. Furthermore, the downregulation of pathways linked to extracellular matrix organization, and proteolysis highlights potential disruptions in microglial structural remodeling. The observed upregulation of PGK1, a key glycolytic enzyme associated with pro-inflammatory microglia, further supports a shift towards an altered metabolic state in *P2ry12* deficient microglia.

### WEIGHTED GENE CO-EXPRESSION NETWORK ANALYSIS (WGCNA) IN WT AND KO MICROGLIA REFLECT FURTHER IMMUNO-METABOLIC PROCESSES

Following our comprehensive DEG analysis of the *P2ry12* deficient microglial expression profile above, we aimed to investigate gene networks disrupted by a genetic deficiency of *P2ry12* and their potential functional associations. To achieve this, we performed Weighted Gene Co-expression Network Analysis (WGCNA), a method used to identify co-expressed gene modules within high-dimensional gene expression data(Langfelder & Horvath, 2008). We constructed a gene co-expression network and proceeded to identify clusters of densely interconnected genes or modules by hierarchically clustering the genes based on the correlation of their expression profiles. The network and the identified modules are depicted in a gene dendrogram obtained by average linkage hierarchical clustering **(Fig 3a)**.

**Figure 3.**
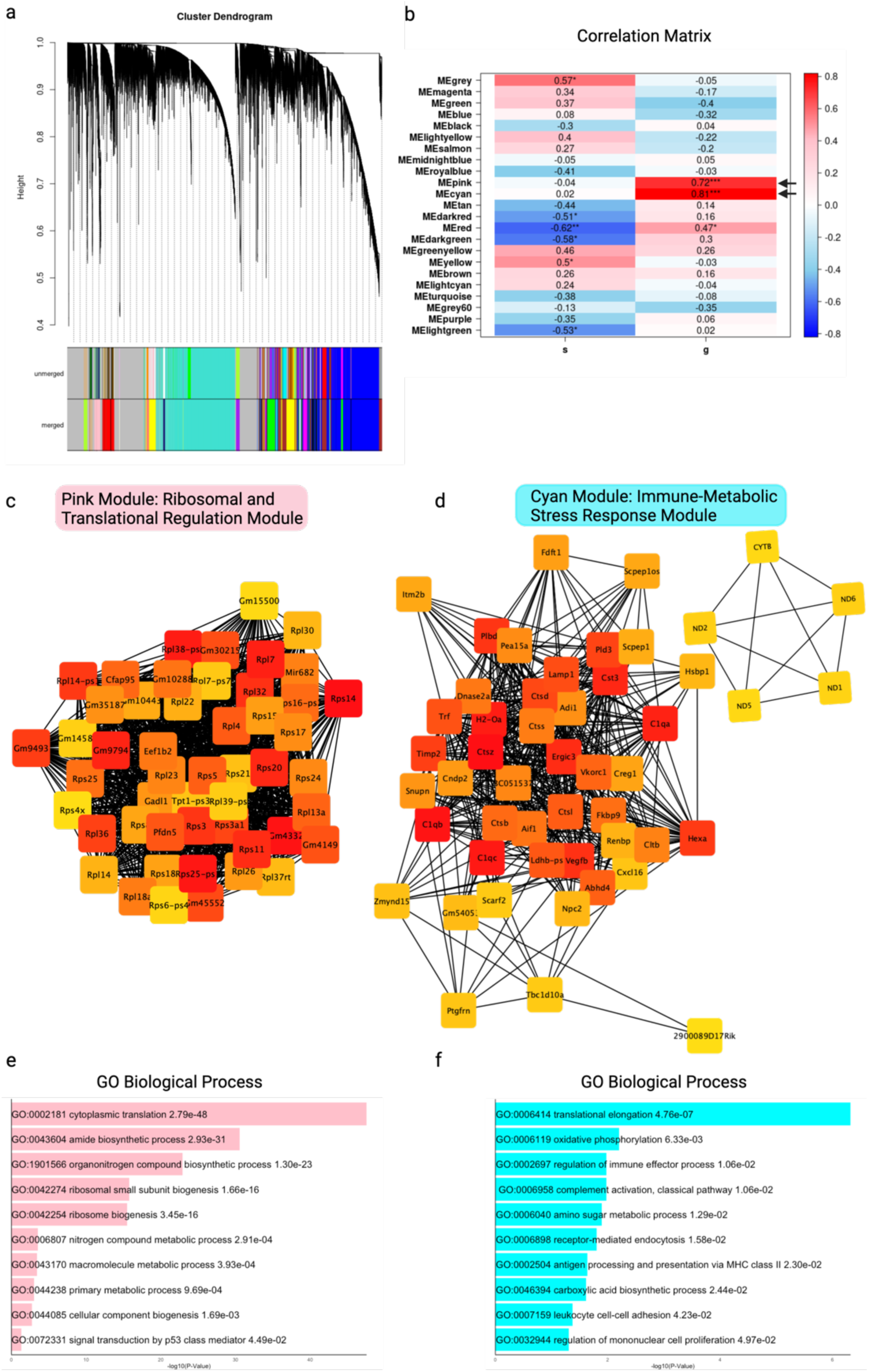
Weighted Gene Co-expression Network Analysis (WGCNA) in WT and KO microglia reflect further immuno-metabolic processes. **(a)** Cluster dendrogram showing co-expressed gene modules identified by WGCNA. Each color represents a different module of co-expressed genes. **(b)** Module-trait correlation heatmap. The pink and cyan modules are significantly correlated with the KO genotype (P < 0.05), indicating potential roles in microglial activation and metabolism. **(c)** Pink Module: Ribosomal and Translational Regulation Module. The network visualization highlights genes involved in ribosome biogenesis and protein synthesis, with hub genes such as Rpl14, Rps21, and Rps3a1. **(d)** GO biological pathways enrichment analysis of the pink module, showing enrichment for processes related to translation, ribosome biogenesis, and macromolecule biosynthesis. **(e)** Cyan Module: Immune-Metabolic Stress Response Module. This network visualization includes genes related to immune activation, oxidative stress, and metabolic regulation. **(f)** GO biological pathways enrichment analysis of the cyan module, showing enrichment for processes such as oxidative phosphorylation, immune regulation, antigen presentation, and leukocyte adhesion. These findings suggest that the cyan module captures genes involved in microglial responses to metabolic stress and inflammation. Data presented as variance-stabilized transformed (VST) counts derived from WGCNA analysis, with visualizations such as cluster dendrograms, module-trait correlation heatmaps, and network graphs to identify co-expression patterns and functional relationships. n = 6 WT (3 male, 3 female) and 10 KO (5 male, 5 female) mice. A soft-thresholding power was selected to approximate scale-free topology and optimize network construction. Hierarchical clustering and dynamic tree cutting were applied to define gene co-expression modules, followed by module eigengene correlation analysis to identify associations between gene modules and experimental traits. The Benjamini-Hochberg correction was used to control false discovery rates in module-trait relationships. Intramodular connectivity and hub gene analysis were performed to determine key regulatory genes within significant modules. Finally, Gene Ontology (GO) enrichment analysis was conducted to functionally annotate gene modules and infer their biological significance.

To assess the relationship between identified modules and genotype, we generated a correlation heatmap. Two modules, the pink and cyan modules, were significantly correlated with the *P2ry12* deficient condition (Pink: r = 0.72, Cyan: r = 0.81) (**Fig 3b**). The pink module (**Fig 3c**) is enriched for genes associated with ribosomal function and protein synthesis, including several ribosomal proteins such as *Rps21*, *Rpl35* and *Rpl13a* (Miller, MacDonald, Kellogg, Karamysheva, & Karamyshev, 2023). The network structure highlights a strong interconnectivity among these genes, suggesting a coordinated activity of translational processes in P2RY12 deficient microglia. GO enrichment analysis of the pink module (**Fig 3e, Supplemental Table 3**) confirmed significant enrichment for biological processes related to translation and ribosomal biogenesis, including cytoplasmic translation (GO:0002181), amide biosynthetic process (GO:0043604), and ribosomal small subunit biogenesis (GO:0042274). These findings indicate that *P2ry12* deficient microglia exhibit enhanced protein synthesis machinery, potentially reflecting increased cellular stress responses.

The cyan module (**Fig 3d**) consists of genes involved in immune activation, lysosomal function, and metabolic stress responses. This module is highly enriched in a cathepsin family genes such as *Ctsb*, *Ctsd*, and *Ctsl*, which are known to regulate lysosomal degradation(Iwama et al., 2021; Nishioku et al., 2002), antigen processing(Clark, Ogbonna, & Malcangio, 2013; K. Zhao, Sun, Zhong, & Luo, 2024), and microglia immune activation(Q. Liu et al., 2019). The strong interconnectivity among these genes suggests that KO microglia undergo transcriptional reprogramming towards an immune-responsive and lysosomal-active state. Interestingly, a previous work showed that loss of *P2ry12* leads to lysosomal accumulation of autofluorescence and neuronal material in microglial soma, suggesting lysosomal dysfunction in *P2ry12* deficient microglia(Bollinger et al., 2023).

Within the cyan module, a distinct subcluster of mitochondrial genes was identified, including several genes encoding for NADH dehydrogenases, which are subunits of mitochondrial complex I (NADH: ubiquinone oxidoreductase), the first enzyme in the electron transport chain and plays a critical role in OXPHOS and ATP production(Sharma, Lu, & Bai, 2009). Alongside *mt-Nd1-6*, the Cytochrome b (*mt-Cytb*) gene is part of mitochondrial Complex III (cytochrome bc1 complex) and plays a crucial role in electron transfer and ATP production(Blakely et al., 2005). GO enrichment analysis of the cyan module confirmed that these genes are involved in immune function, antigen processing, and metabolic stress responses, including translational elongation (GO:0064114), oxidative phosphorylation (GO:006119), antigen processing and presentation via MHC class II (GO:002509), and receptor-mediated endocytosis (GO:0006898) (**Fig. 3f, Supplemental Table 4**). The cyan module, enriched in mitochondrial and immune-related genes, suggests that *P2ry12*-deficient microglia undergo a metabolic shift characterized by elevated mitochondrial activity, increased energy demands, and heightened oxidative stress, potentially influencing their immune and neuroinflammatory responses.

### A *P2ry12* DEFICIENCY IMPAIRS THE MICROGLIAL TRANSCRIPTIONAL RESPONSE TO INFLAMMATORY STIMULI

To investigate how a *P2ry12* deficiency affects microglia under pathological conditions, we turned to a functional assessment and examined microglial responses to inflammatory stimuli. WT and *P2ry12* deficient littermate mice were injected with LPS (1 mg/kg; i.p.) followed by an evaluation of the microglial transcriptional state 24 h after injection (**Fig 4a**).

**Figure 4.**
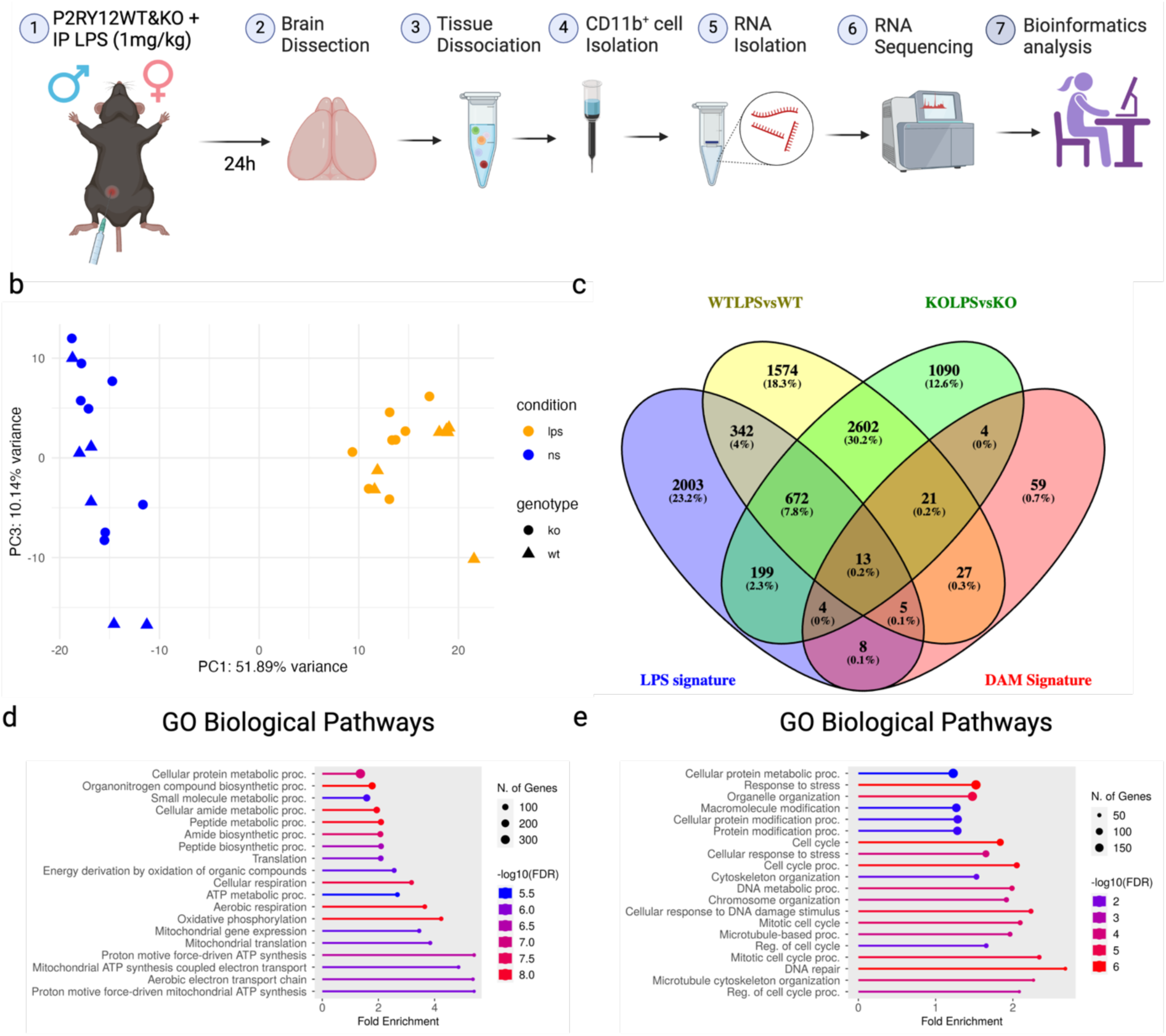
A *P2ry12* deficiency impairs microglial reactivity to inflammatory stimuli. **(a)** Schematic depicting the experimental approach. WT and KO mice were treated with LPS (1 mg/kg) or left untreated (NS, no stimulation). Brain tissue was collected at 24h post-injection for microglia isolation, RNA extraction, bulk RNA sequencing, and bioinformatics analysis. **(b)** PCA plot between groups based on genotype (WT vs. KO) and treatment (LPS vs. NS). **(c)** Venn diagram comparing WT-LPS and KO-LPS DEGs with DAM DEGs as well as LPS-activated microglia DEGs described by (Sabogal-Guáqueta et al., 2023). **(d)** GO biological pathways enrichment analysis for the 1574 DEGs exclusively in the WT-LPS showing enrichment of protein synthesis and oxidative phosphorylation related terms. **(e)** GO biological pathways enrichment analysis for the 1090 DEGs exclusively in the KO-LPS highlights associations with cell cycle regulation, DNA maintenance, and cellular stress responses. Data presented as variance-stabilized transformed (VST) counts with visualizations such as PCA plots to accurately depict expression distributions. n = 13 WT (6 male and 7 female) and 16 KO (8 male and 8 female) mice. The following tests were performed for statistical analysis: Negative binomial GLM with the Wald test for differential expression, Benjamini-Hochberg correction to control false discoveries, apeglm shrinkage to reduce variability in LFC estimates and improve biological interpretability.

Then, we performed PCA on RNAseq data from microglia to determine how genotype and LPS treatment influence transcriptional variability (**Fig 4b**). PC1 (52% variance) largely separates samples based on LPS treatment, indicating a strong transcriptional shift between non-stimulated (NS) and LPS-treated microglia. PC3 (10% variance) captures additional variability, but there is no strong separation by genotype (WT vs. KO) within each treatment condition. However, most of the LPS-activated KO microglia (KO-LPS) samples slightly separate from the LPS-activated WT microglia (WT-LPS) samples. This suggests that LPS exposure is the dominant factor driving transcriptional changes in microglia, while a *P2ry12* deficiency, though affecting the microglial transcriptional response (as discussed below), is a less dominant driver under these conditions.

To identify genes differentially regulated by a *P2ry12* deficiency in response to LPS, we performed DEG analysis comparing KO-LPS with untreated KO microglia (KO-LPS vs KO), as well as WT-LPS with untreated WT microglia (WTLPS vs WT). Next, using a Venn diagram we compared the obtained DEG lists to the DAM signature, as well as with the DEGs previously described for LPS-activated mouse microglia(Sabogal-Guáqueta et al., 2023). We found that DEGs resulting from WT-LPS vs WT and KO-LPS vs KO shared more genes with the LPS signature (343 and 199 genes) than with the DAM signature (27 and 4 genes). In this context, our WTLPS vs WT DEGs overlapped more with the previously reported LPS and DAM signatures than our KOLPS vs KO DEGs (**Fig 4c**). From the Venn diagram we identified 1574 DEGs exclusively in the WTLPS vs WT and 1090 DEGs in the KOLPS vs KO. We took these 2 lists of genes and performed GO biological pathways enrichment analysis which revealed the enrichment of pathways associated with protein metabolism and oxidative phosphorylation in the WT microglia when treated with LPS (**Fig 4d, Supplemental Table 5**). Conversely, we found that the unique DEGs in the LPS-treated KO microglia enriched pathways related to cell cycle and DNA damage responses (**Fig 4e, Supplemental Table 6**). In the cell cycle associated genes we found important protein kinases such as *Chk1*, *Chk2* and *Wee1*(**Supplemental Table 6**) that prevent cell cycle progression upon DNA damage (Drew, Zenke, & Curtin, 2025), that were uniquely upregulated in the LPS-KO microglia.

### A *P2ry12* DEFICIENCY ALTERS MICROGLIAL TRANSCRIPTIONAL PROFILE TOWARD INCREASED FERROPTOSIS SUSCEPTIBILITY FOLLOWING LPS-INDUCED INFLAMMATION

We kept examining expression patterns between the untreated and LPS-treated conditions and found two interesting group of genes (**Fig 5a**): group 1 included genes that are upregulated in the KO cells with LPS treatment but not affected by LPS in the WT cells (top) including *Nrgn*, *Rxrg* and *Apob*. Genes in group 2 were upregulated in the WT when treated with LPS but remained unchanged or unresponsive in the KO under LPS (bottom): including *Ccr1, Traf1* and *Slc7a11*.

**Figure 5.**
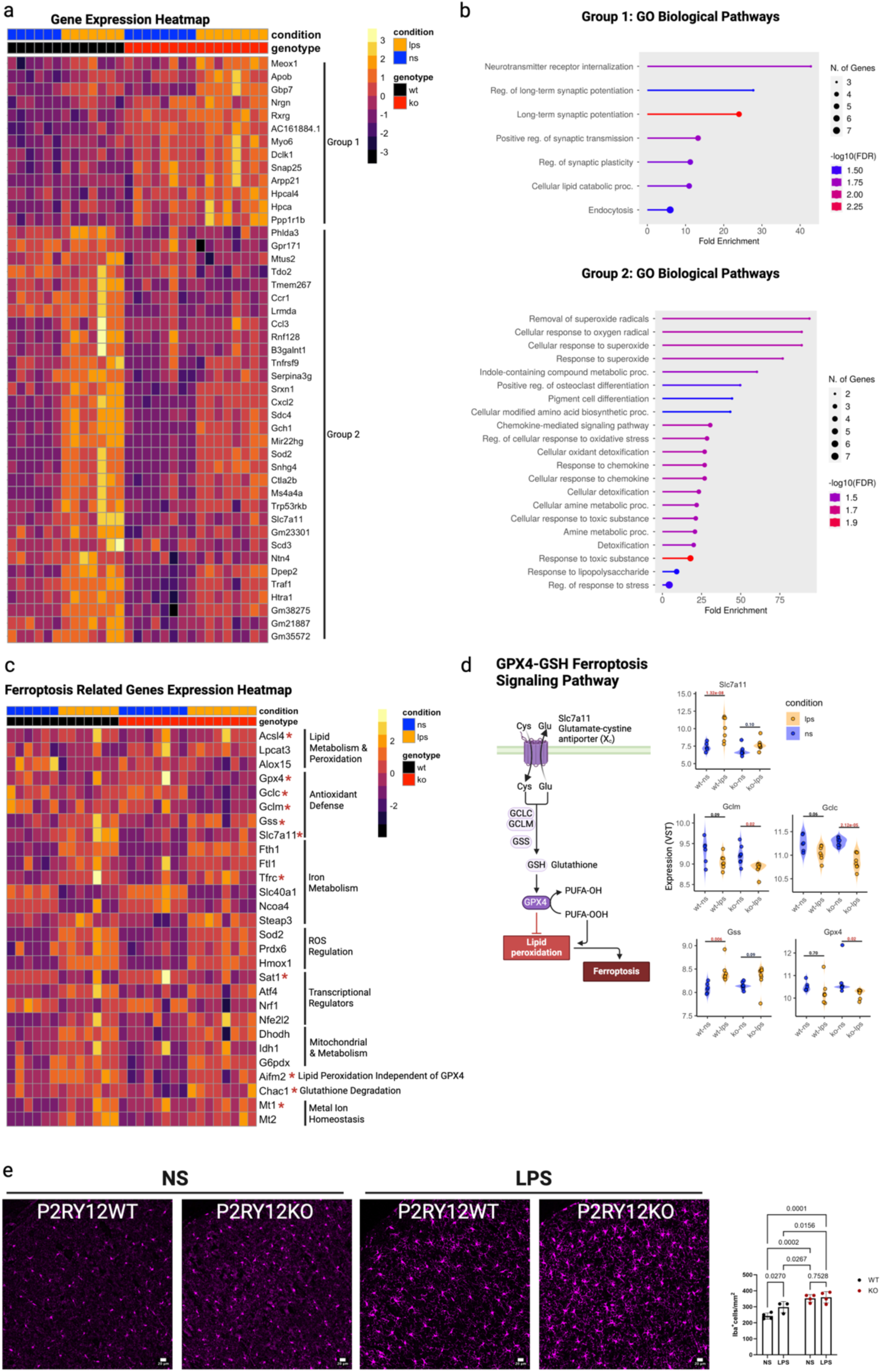
*P2ry12* deficiency enhances microglial vulnerability to ferroptosis in response to LPS-induced inflammation. **(a)** Heatmap of expression patterns between the WT, KO, WT-LPS, KO-LPS conditions with genes that are upregulated in the KO but not affected by LPS in the WT (top) and genes upregulated in the WT-LPS but unresponsive in the KO-LPS (bottom). **(b)** Genes upregulated in the KO but not affected by LPS in the WT (Group 1, top) and GO enrichment analysis of genes upregulated in the WT-LPS but unresponsive in the KO-LPS (Group 2, bottom). **(c)** Heatmap depicting the expression of ferroptosis-associated genes across different conditions (WT-NS, WT-LPS, KO-NS, KO-LPS). **(d)** Left: Schematic representation of the GPX4-GSH antioxidant pathway, showing key enzymes involved in ferroptosis regulation, including SLC7A11, GCLC, GCLM, GSS, and GPX4. Right: Scatter plots of transcript levels for ferroptosis-related genes across conditions (LPS vs. NS). Red p-values indicate statistically significant differences between conditions. Data presented as log-transformed normalized counts with visualizations such as PCA plots, heatmaps and scatter plots to accurately depict expression distributions. n = 13 WT (6 male and 7 female) and 16 KO (8 male and 8 female) mice. The following tests were performed for statistical analysis: Negative binomial GLM with the Wald test for differential expression, Benjamini-Hochberg correction to control false discoveries, apeglm shrinkage to reduce variability in LFC estimates and improve biological interpretability. **(e)** Left: Immunofluorescence staining of microglia (IBA1, magenta) across non stimulated (NS) and LPS treated WT and KO littermates brains. Right: The quantification of IBA1 positive cells shows a significant increase in WT microglia after LPS treatment (P = 0.0270), however KO microglia doesn’t exhibit any increase. Data presented as mean ± s.e.m. n = 7 female WT (4 NS and 3 LPS treated) and 8 female KO (4 NS and 4 LPS treated) mice. Data analysis was performed with 2-way ANOVA to assess the effects of both genotype and treatment and presented as mean ± SEM. Tukey’s multiple comparisons test was applied for post hoc analysis. A p-value < 0.05 was considered significant.

GO enrichment analysis of genes in group 1 revealed several pathways related to synaptic modulation, lipid metabolism, and endocytosis (**Fig 5b, top**) (**Supplemental Table 7**). Specifically, we observed enrichment for long-term synaptic potentiation, suggesting potential changes in synaptic plasticity and neuronal communication. Additionally, the lipid catabolic process was significantly upregulated, indicating altered lipid metabolism, which may influence further microglial energy homeostasis and inflammatory responses. Other enriched pathways included regulation of endocytosis highlighting potential changes in cellular uptake mechanisms relevant to antigen processing and immune function.

Next, we performed GO enrichment analysis of the unresponsive genes in group 2 (**Fig 5b, bottom**) (**Supplemental Table 7**) which highlights immune activation and oxidative stress responses. Notably, we observed enrichment for cellular detoxification and response to oxidative stress, suggesting that *P2ry12* deficient microglia may exhibit increased susceptibility to oxidative damage under LPS stimuli. Several immune-related pathways were also significantly enriched, highlighting altered immune reactivity in the *P2ry12* deficient microglia when exposed to LPS. Specifically, response to chemokine and chemokine-mediated signaling suggest increased microglial communication and an overall heightened inflammatory response.

One of the significantly downregulated genes in KO microglia compared to WT-LPS was *Slc7a11* (**Fig 5a**). This gene encodes for the cystine/glutamate antiporter protein known as xCT−(Lupica-Tondo, Arner, Mogilenko, & Voss, 2024) which when combined with SLC3A2 form system x_c-_. In this sense, system x_c-_is crucial for cysteine import for the cell, which is the substrate for glutathione synthesis and a key regulator of ferroptosis resistance. Given this finding, we evaluated the expression of other ferroptosis-associated genes to determine whether *P2ry12* deficiency increases susceptibility to ferroptosis following LPS stimulation.

One of the most striking findings from the ferroptosis-related gene expression heatmap (**Fig 5c**) was the widespread dysregulation of antioxidant defense genes in P2RY12 deficient microglia following LPS stimulation. Several genes critical for Glutathione (GSH) biosynthesis and antioxidant defense were significantly impacted in KO-LPS microglia (**Fig 5d**), including: *Slc7a11* (cystine/glutamate exchanger) that mediates cystine uptake into cells, *Gclc* (Glutamate-Cysteine Ligase Catalytic Subunit) a rate-limiting enzyme in glutathione synthesis, *Gclm* (Glutamate-Cysteine Ligase Modifier Subunit), a regulatory subunit that enhances *Gclc* activity, *Gss* (Glutathione Synthetase) which catalyzes the final step in glutathione biosynthesis, and *Gpx4* (Glutathione Peroxidase 4) which uses glutathione to detoxify lipid hydroperoxides, preventing ferroptosis(X. Yu & Long, 2016). These results suggest a profound impairment in glutathione homeostasis in KO-LPS microglia, that one could predict would lead to increased vulnerability to oxidative stress and ferroptosis. Based on this observation we sought to quantify microglia numbers across conditions (**Fig 5e**). We found an increase in microglia (IBA1^+^) positive cells in the somatosensory cortex of LPS treated WT mice. However, this was not the case for the KO brains, where microglia numbers remained stable, suggesting that number changes elicited by LPS in the WT are not recapitulated by *P2ry12* deficient microglia (**Fig 5e**).

## Discussion

As both immune and brain-resident cells, microglia exhibit a uniquely dynamic nature, making their activation states difficult to rigidly classify(Paolicelli et al., 2022). A key aspect of this uniqueness is the high expression of *P2ry12* by microglia in the brain, which is often downregulated in response to changes in the brain microenvironment as occurs in pathology(Bohlen et al., 2017; A. Deczkowska et al., 2018). Notably, *P2ry12* expression diminishes when microglia are removed from their native CNS context, as observed when they are isolated from the brain(Bohlen et al., 2017; Cadiz et al., 2022). This presents a significant limitation in studying the receptor’s function *in vitro*. Therefore, in this study, we aimed to investigate the role of P2RY12 in the microglial homeostatic state and modulating responses to inflammatory stimuli by examining the transcriptional profile of *P2ry12* deficient microglia compared to *P2ry12*-sufficient littermates. We show that *P2ry12* deficient microglia retained their transcriptional identity with the upregulation of a few recognized DAM signature genes such as: *B2m, Ctsb* and *Fth1* (**Fig 1**). Additionally, *P2ry12* deficient microglia exhibited perturbations in genes associated with cellular metabolism and immune signaling (**Fig 2**). Furthermore, we observed similar disturbances within regulatory pathways (**Fig 3**) indicating that *P2ry12* deficient microglia are in a more metabolically active state that could be compatible with a metabolic reprogramming phase in preparations to transition to an activated state. Upon LPS challenge, microglia showed unique DEGs involved in protein metabolism and oxidative phosphorylation in the WT microglia, whereas the unique KO DEGs are associated with DNA damage response and oxidative stress (**Fig 4**). Additionally, LPS-treated KO microglia exhibited an “unresponsive” set of genes associated with immune activation and oxidative stress responses suggesting that these processes are, at least, partly under P2RY12 control in microglia. Strikingly, we found impaired GPX4-Glutathione (GSH) antioxidant responses in the LPS-stimulated KO microglia, rendering them likely more susceptible to ferroptotic insults (**Fig 5**). Our findings provide novel evidence that P2RY12 orchestrate microglial immunometabolic programming and activation especially to inflammatory stimuli.

Due to *P2ry12*’s expression sensitivity during microglial isolation, its role in microglial function remains poorly understood as it can largely only be studies in microglia *in vivo* and not in culture since it is significantly downregulated in culture(Bohlen et al., 2017). However, this is not the case for platelets, the other primary site of *P2ry12* expression in the body where P2RY12 has been described to have a constitutive role associated with the inhibition of adenylate cyclase thus reducing levels of cyclic AMP. This constitutive role in platelets was shown to be essential for priming platelets to evoke a more rapid platelet aggregation response upon P2RY12-ligand activation(Garcia et al., 2019). In this regard, our study provides evidence that P2RY12 might also have a constitutive function in microglia that involves intracellular signaling regulation as evidenced by the fact that with a *P2ry12* deficiency, microglia exhibited an expression profile consistent with a highly metabolic state with increased cellular energy production through both glycolysis (**Fig 2a, d, e**) and OXPHOS (**Fig 2d and Fig 3e-f**), as well as protein biosynthesis (**Fig 2c-d and Fig 3c-d**).

Interestingly, our results show that *P2ry12* deficient microglia upregulate the expression of the dimethylarginine dimethylaminohydrolase 1 (*ddha1*) enzyme that acts to degrade ADMA, an intracellular inhibitor of nitric oxide synthase (NOS). Hence, an upregulation of *ddha1* implies an increased production of NO (Reddy et al., 2018). Since NO has known vasodilatory effects, these results align with our previous findings in *P2ry12* deficient mice that showed increased cerebral blood flow and stunted hypercapnic response(Bisht et al., 2021). Conversely, the downregulated DEGs in P2RY12 deficient microglia included genes associated with ECM turnover and remodeling (**Fig 2b**). Impaired ECM remodeling can have an impact on vascular homeostasis which can influence blood flow by promoting ECM-components deposition along the vasculature and ECM disorganization(Chauhan et al., 2008; Delhon et al., 2019; Gandhi, Khan, Lentz, & Chauhan, 2012; Le Goff et al., 2008; Lin et al., 2024). This is a possibility that should be further investigated.

A recent proteomic study examining the brains of mice lacking the key microglial-signaling receptors P2RY12 and CX3CR1(Strohm et al., 2025) identified proteins highly correlated with the P2RY12-deficient genotype, linking them to acetyl-CoA metabolism and protein biosynthesis-related processes. While this study differs methodologically from ours—particularly in microglial specificity (our study focuses on isolated microglia, whereas that proteomic study analyzed whole-brain lysates) and in the use of translational and transcriptional inhibitors to delay microglial activation during brain processing—both investigations converge on similar conclusions that microglial (and brain metabolic states) are altered in *P2ry12*-deficient cells and brains. These findings strongly support the notion that P2RY12 plays a basal role in regulating microglial metabolism, ensuring their ability to meet cellular demands in response to their physiological or pathological environment. Together, that study(Strohm et al., 2025) and this one suggest that metabolic effects of a *P2ry12* deficiency may extend beyond microglia to the whole brain.

Under high energy demanding states or oxidative stress, cells generate reactive oxygen species (ROS). We show that in the basal state, P2RY12 deficient microglia seem to be in a high ATP producing state that might be leading to an increase in ROS generation and oxidative stress, suggested by increased expression levels of metallothioneins 1 and 2 (*Mt1* and *Mt2)* and ferritin (*Fth1 and Ftl*) (**Fig 2d**). These proteins regulate metal homeostasis and ferric iron binding (**Fig 2c, Supplemental Table 2**), with a key role in preventing free-radical damage and antioxidant response(Atrián-Blasco et al., 2017; Klaassen, Liu, & Choudhuri, 1999; Lazo & Pitt, 1995; G. Zhao, Arosio, & Chasteen, 2006). The increase in expression of these genes in *P2ry12* deficient microglia might be counteracting cellular damage under the highly demanding metabolic state. Of particular interest ferritin heavy and light chain expression is regulated by iron levels(W. Li et al., 2015), the increase in *fth1* and *ftl* expression in the KO microglia (**Fig 1c,e and Fig 2d**) might also suggest changes in iron metabolism.

Our results show that *P2ry12* deficient microglia have exacerbated deficits in the antioxidant response upon LPS challenge (**Fig 5a, 5c**). Transcriptionally, several genes related to GSH production are dysregulated in *P2ry12* deficient microglia, including a failed upregulation of *Slc7a11* after LPS treatment (**Fig 5a-d**). Strikingly, *Slc7a11* is a direct target gene suppressed by p53 an important mediator of ferroptosis susceptibility(Jiang et al., 2015; Yanqing Liu & Gu, 2022). According to our study, in the basal state, KO microglia exhibit the upregulation of *Rps15*, *Rps20* and *Rpl37a* genes that inhibit MD2/MD4 function (**Fig 2b and Fig 3c, Supplemental Table 1**). MD2/MD4 serve as a negative regulator of p53 DNA binding activity(Daftuar et al., 2013).

Additionally, upon LPS stimulation, KO microglia uniquely upregulate genes involved in cell cycle checkpoint control, including *Chek1*, *Chek2*, and *Wee1* (**Fig. 4e**; **Supplemental Table 6**). These genes are central regulators of the DNA damage response (DDR), with CHK1/2 activation notably linked to p53-dependent growth arrest(Fagagna et al., 2003; Gire, Roux, Wynford-Thomas, Brondello, & Dulic, 2004)

We propose that the highly metabolically active basal state of KO microglia may prime them for oxidative stress. When challenged with LPS, this oxidative burden may surpass their antioxidant capacity, leading to DNA damage and subsequent induction of *Chek1/2* and *Wee1* expression. Oxidative stress has also been shown to stabilize p53 via post-translational mechanisms(Kruse & Gu, 2009), which could further suppress *slc7a11* expression, a gene repressed by p53, thereby impairing glutathione (GSH)-mediated antioxidant defenses.

Together, these findings suggest that in *P2ry12*-deficient microglia, a defective antioxidant response, coupled with altered energy and iron metabolism, may increase susceptibility to ferroptosis during microglia activation. Notably, while LPS-treated WT microglia show increased IBA1⁺ cell numbers, suggesting a proliferative response, KO microglia do not (**Fig. 5e**).

Ferroptosis is a form of regulated necrotic cell death characterized by the iron-dependent accumulation of oxidized phospholipids that compromise cell membrane(Dixon et al., 2012; Stockwell et al., 2017). Under homeostatic conditions, ferroptosis is inhibited primarily by GSH-GPX4 mediated antioxidant response(Seibt, Proneth, & Conrad, 2019). Recent research has implicated Aberrant ferroptosis related gene expression to CNS pathologies like AD(H. Zhao et al., 2023) and epilepsy(X. Li et al., 2024), especially in macrophages and microglia, highlighting the intricate relationship between ferroptosis, oxidative stress and neuroinflammation. Here, macrophages exhibit iron metabolism alterations and differences in ferroptosis sensitivity depending on their activation state(Lupica-Tondo et al., 2024; Yang et al., 2022). This is especially relevant since P2RY12 is downregulated in human and mouse AD models(A. Deczkowska et al., 2018; Kenkhuis et al., 2022; H. Keren-Shaul et al., 2017), where ferroptosis is known to occur. Interestingly, ferroptosis can propagate among cells in a wave-like manner preceding cell death(Kim et al., 2016; Linkermann et al., 2014; Riegman et al., 2020). While this is a relatively new area of investigation, emerging evidence suggests that ferroptosis is not a purely cell-autonomous process. Specifically, studies have shown that microglia can act as initiators of a ferroptotic cascade, with astrocytes serving as intermediaries that ultimately lead to neuronal death (Liddell et al., 2024). This multicellular propagation of ferroptotic stress highlights the potential for microglial vulnerability to drive broader network-level dysfunction.

Our findings suggest that P2RY12-deficient microglia exhibit transcriptional features consistent with heightened ferroptosis susceptibility. In conditions where *P2ry12* is persistently downregulated, such as in AD where microglia with low *P2ry12* expression cluster around amyloid-beta plaques(Walker et al., 2020), or in neurological disorders like epilepsy where microglial P2RY12 may contribute to seizure severity(Eyo et al., 2014; Q. Wang et al., 2023), this vulnerability may compromise microglial neuroprotective roles, especially when further challenged, such as in systemic infections, documented to worsen these pathologies (Holmes et al., 2009; Perry, Newman, & Cunningham, 2003; Vezzani et al., 2016).

It has been shown that a subset of microglia is able to engage in long-term iron storage and these cells are especially prone dystrophy in the aged brain, suggesting that chronic iron load, diminished microglial antioxidant and self-renewal capacity underlie microglial dystrophy (Lopes, Sparks, & Streit, 2008). In conditions such as AD, where iron-rich amyloid plaques and chronic inflammation converge, this subpopulation may be particularly vulnerable to ferroptosis. Thus, in the absence of *P2ry12*, microglia may become sources of oxidative damage due to dysfunctional iron handling, accelerating iron-mediated pathology. Moreover, since ferroptosis is increasingly recognized as a key contributor to the pathophysiology of neurodegenerative and neurological disorders, investigating the intersection of P2RY12 signaling, redox regulation, and ferroptotic vulnerability in microglia offers a promising avenue for uncovering novel mechanisms of disease progression.

Finally, there are well-documented differences between human and mouse microglia, which pose important considerations when translating findings from mouse models to human neurobiology and disease. However, P2RY12, expression is conserved between human and mouse microglia under homeostatic conditions(Geirsdottir et al., 2019; Gosselin et al., 2017; Hasselmann et al., 2019). Moreover, its downregulation has been corroborated in human microglia in the context of neuroinflammation and aging (Masuda et al., 2019; Palmer, Liu, Romanow, Lee, & Chun, 2021; Škandík et al., 2025), supporting the translational relevance of this study. Our finding that P2RY12 loss disrupts microglial redox balance and increases vulnerability to ferroptosis coupled with evidence of impaired redox homeostasis and oxidative injury in human neurodegenerative diseases, highlights a critical and potentially conserved function of P2RY12 signaling in microglial resilience. Additionally, since P2RY12 is a known target of antiplatelet therapies used clinically, and such agents could cross into the CNS under conditions of blood-brain barrier (BBB) compromise, these findings raise important questions about the possibility of unintended effects of systemic medications on microglial health and function in patients with neuroinflammatory conditions.

## Limitations of this Study

This study has several limitations that warrant consideration. First, the use of a global P2ry12 knockout (KO) model may introduce systemic effects, such as potential coagulation abnormalities, that could influence iron metabolism and confound microglia-specific interpretations. To better isolate the direct effects of P2ry12 loss in microglia from broader systemic alterations, future studies using conditional, microglia-specific P2ry12 knockout models will be essential.

Second, this study relies primarily on transcriptomic and bioinformatic analyses, which are inherently hypothesis-generating. While our data suggest that P2RY12 deficiency alters microglial redox balance, immune activation, and ferroptosis susceptibility, future studies will be necessary to directly test these predicted phenotypes in disease-relevant models and determine their impact on CNS function. Additionally, the transcriptomic analyses were performed on RNA extracted from whole-brain lysates, which may obscure region-specific microglial transcriptional differences particularly in brain regions more vulnerable to inflammatory stimuli such as the hippocampus (Jung et al., 2023). Future studies employing region-specific isolation strategies will be important to reveal spatial heterogeneity in microglial responses.

Third, although we confirmed the upregulation of PGK1 at the protein level (**Fig. 2d**), additional functional metabolic assays are required to comprehensively characterize metabolic reprogramming in P2RY12-deficient microglia beyond transcriptional and proteomic data. While technically challenging, especially in vivo, such assays would strengthen the mechanistic interpretation of the metabolic phenotype observed.

Fourth, the systemic LPS model used in this study induces microglial activation without extensive tissue damage. We acknowledge that models with more overt neurodegeneration, such as experimental autoimmune encephalomyelitis (EAE), or models of epilepsy and AD, would be additionally suited to reveal in vivo functional consequences of P2RY12 deficiency in such degenerative contexts. However, our goal was to characterize transcriptional and metabolic changes in microglia under a sublethal inflammatory challenge. This approach allowed us to uncover early immuno-metabolic vulnerabilities without the confounding influence of tissue damage. While we focused on microglial responses, we recognize the importance of examining broader CNS phenotypes, including potential effects on neurons and other glia. We note that future studies using cell type-specific models will be critical to disentangle direct versus secondary effects of P2RY12 deficiency including in more severe neuroinflammatory models to further validate and extend our findings in disease-relevant contexts.

Finally, while our study included both male and female mice, in some cases the sample size per sex was limited to n=3 and prevents definitive conclusions regarding sex differences in P2RY12 deficient microglia.

Despite these limitations, our findings provide novel insights into the role of P2RY12 in regulating microglial immune-metabolic states and ferroptosis susceptibility. These observations lay the groundwork for future investigations into region- and sex-specific microglial-P2RY12 dynamics in neurodegenerative and neuroinflammatory conditions.

## Gene Abbreviation List

*Adamts13*: A Disintegrin And Metalloproteinase With Thrombospondin Motifs 13
*Adamts16*: A Disintegrin And Metalloproteinase With Thrombospondin Motifs 16
*Adamtsl2*: ADAMTS Like 2
*Apob*: Apolipoprotein B
*B2m*: Beta-2-Microglobulin
*Ccr1*: C-C Motif Chemokine Receptor 1
*Ccrl2*: C-C Motif Chemokine Receptor Like 2
*Cst7*: Cystatin F
*Ctsb*: Cathepsin B
*Ddah1*: Dimethylarginine Dimethylaminohydrolase 1
*Ddx3y*: DEAD-Box Helicase 3 Y-Linked
*Eif2s3y*: Eukaryotic Translation Initiation Factor 2 Subunit 3 Y-Linked
*Fth1*: Ferritin Heavy Chain 1
*Ftl*: Ferritin Light Chain
*Gclc*: Glutamate-Cysteine Ligase Catalytic Subunit
*Gclm*: Glutamate-Cysteine Ligase Modifier Subunit
*Gpx4*: Glutathione Peroxidase 4
*Gss*: Glutathione Synthetase
*Gsto1*: Glutathione S-Transferase Omega 1
*Ighv1-18*: Immunoglobulin Heavy Variable 1-18
*Igkv19-93*: Immunoglobulin Kappa Variable 19-93
*Kdm5d*: Lysine Demethylase 5D
*Lox*: Lysyl Oxidase
*Med12l*: Mediator Complex Subunit 12 Like
*Mt1*: Metallothionein 1
*Mt2*: Metallothionein 2
*mt-Cox1*: Cytochrome c oxidase subunit I
*mt-Cox7a2l*: Cytochrome c Oxidase Subunit VIIa Polypeptide 2 Like
*mt-Cytb*: Cytochrome b
*mt-Nd1*: NADH:Ubiquinone Oxidoreductase Core Subunit 1
*mt-Nd2*: NADH:Ubiquinone Oxidoreductase Core Subunit 2
*mt-Nd3*: NADH:Ubiquinone Oxidoreductase Core Subunit 3
*mt-Nd4*: NADH:Ubiquinone Oxidoreductase Core Subunit 4
*mt-Nd5*: NADH:Ubiquinone Oxidoreductase Core Subunit 5
*mt-Nd6*: NADH:Ubiquinone Oxidoreductase Core Subunit 6
*Nfkbia*: Nuclear Factor Of Kappa Light Polypeptide Gene Enhancer In B-Cells Inhibitor, Alpha
*Nrgn*: Neurogranin
*Pgk1*: Phosphoglycerate Kinase 1
*Rxrg*: Retinoid X Receptor Gamma
*Slc7a11*: Solute Carrier Family 7 Member 11
*Traf1*: TNF Receptor Associated Factor 1
*Tyrobp*: Transmembrane Immune Signaling Adaptor TYROBP
*Uty*: Ubiquitously Transcribed Tetratricopeptide Repeat Containing, Y-Linked
*Xist*: X Inactive Specific Transcript

## References

Allen, M., Zou, F., Chai, H. S., Younkin, C. S., Miles, R., Nair, A. A., … Rowley, C. N. (2012). Glutathione S-transferase omega genes in Alzheimer and Parkinson disease risk, age-at-diagnosis and brain gene expression: an association study with mechanistic implications. Molecular neurodegeneration, 7, 1–12.

Atrián-Blasco, E., Santoro, A., Pountney, D. L., Meloni, G., Hureau, C., & Faller, P. (2017). Chemistry of mammalian metallothioneins and their interaction with amyloidogenic peptides and proteins. Chemical Society Reviews, 46(24), 7683–7693.

Badimon, A., Strasburger, H. J., Ayata, P., Chen, X., Nair, A., Ikegami, A., … Uweru, J. O. (2020). Negative feedback control of neuronal activity by microglia. Nature, 586(7829), 417–423.

Baik, S. H., Ramanujan, V. K., Becker, C., Fett, S., Underhill, D. M., & Wolf, A. J. (2023). Hexokinase dissociation from mitochondria promotes oligomerization of VDAC that facilitates NLRP3 inflammasome assembly and activation. Science immunology, 8(84), eade7652.

Basilico, B., Ferrucci, L., Ratano, P., Golia, M. T., Grimaldi, A., Rosito, M., … Marrone, M. C. (2022). Microglia control glutamatergic synapses in the adult mouse hippocampus. Glia, 70(1), 173–195.

Bennett, F. C., Bennett, M. L., Yaqoob, F., Mulinyawe, S. B., Grant, G. A., Gephart, M. H., … Barres, B. A. (2018). A combination of ontogeny and CNS environment establishes microglial identity. Neuron, 98(6), 1170–1183. e1178.

Bernier, L.-P., York, E. M., & MacVicar, B. A. (2020). Immunometabolism in the brain: how metabolism shapes microglial function. Trends in Neurosciences, 43(11), 854–869.

Bisht, K., Okojie, K. A., Sharma, K., Lentferink, D. H., Sun, Y. Y., Chen, H. R., … Eyo, U. B. (2021). Capillary-associated microglia regulate vascular structure and function through PANX1-P2RY12 coupling in mice. Nat Commun, 12(1), 5289. doi:10.1038/s41467-021-25590-8

Blakely, E. L., Mitchell, A. L., Fisher, N., Meunier, B., Nijtmans, L. G., Schaefer, A. M., … Taylor, R. W. (2005). A mitochondrial cytochrome b mutation causing severe respiratory chain enzyme deficiency in humans and yeast. The FEBS journal, 272(14), 3583–3592.

Board, P., Chelvanayagam, G., Jermiin, L., Tetlow, N., Tzeng, H.-F., Anders, M., & Blackburn, A. (2001). Identification of novel glutathione transferases and polymorphic variants by expressed sequence tag database analysis. Drug metabolism and disposition, 29(4), 544–547.

Bohlen, C. J., Bennett, F. C., Tucker, A. F., Collins, H. Y., Mulinyawe, S. B., & Barres, B. A. (2017). Diverse Requirements for Microglial Survival, Specification, and Function Revealed by Defined-Medium Cultures. Neuron, 94(4), 759–773 e758. doi:10.1016/j.neuron.2017.04.043

Bollinger, J. L., Dadosky, D. T., Flurer, J. K., Rainer, I. L., Woodburn, S. C., & Wohleb, E. S. (2023). Microglial P2Y12 mediates chronic stress-induced synapse loss in the prefrontal cortex and associated behavioral consequences. Neuropsychopharmacology, 48(9), 1347–1357.

Bou-Abdallah, F., Paliakkara, J. J., Melman, G., & Melman, A. (2018). Reductive mobilization of iron from intact ferritin: mechanisms and physiological implication. Pharmaceuticals, 11(4), 120.

Cadiz, M. P., Jensen, T. D., Sens, J. P., Zhu, K., Song, W.-M., Zhang, B., … Fryer, J. D. (2022). Culture shock: microglial heterogeneity, activation, and disrupted single-cell microglial networks in vitro. Molecular neurodegeneration, 17(1), 26.

Cao, W., Feng, Z., Zhu, D., Li, S., Du, M., Ye, S., … Fang, Y. (2023). The role of PGK1 in promoting ischemia/reperfusion injury-induced microglial M1 polarization and inflammation by regulating glycolysis. NeuroMolecular Medicine, 25(2), 301–311.

Chauhan, A. K., Kisucka, J., Brill, A., Walsh, M. T., Scheiflinger, F., & Wagner, D. D. (2008). ADAMTS13: a new link between thrombosis and inflammation. Journal of Experimental Medicine, 205(9), 2065–2074.

Clark, A. K., Ogbonna, A. C., & Malcangio, M. (2013). Cathepsin and Microglia. In G. F. Gebhart & R. F. Schmidt (Eds.), Encyclopedia of Pain (pp. 472–483). Berlin, Heidelberg: Springer Berlin Heidelberg.

Cordoba, C. M., Barrera, M., Perdomo, S., Franco, P. G., Alvarez, J. A., Dueñas, Z. Y., & Avila, M. Y. (2021). Inhibitor of Lysyl Oxidase in The Optic Nerve Head Complex Imparts Partial Protection Against Injury in Experimental Glaucoma.

Couto, N., Wood, J., & Barber, J. (2016). The role of glutathione reductase and related enzymes on cellular redox homoeostasis network. Free radical biology and medicine, 95, 27–42.

Cronk, J. C., Filiano, A. J., Louveau, A., Marin, I., Marsh, R., Ji, E., … Acton, S. (2018). Peripherally derived macrophages can engraft the brain independent of irradiation and maintain an identity distinct from microglia. Journal of Experimental Medicine, 215(6), 1627–1647.

Császár, E., Lénárt, N., Cserép, C., Környei, Z., Fekete, R., Pósfai, B., … Szabadits, E. (2022). Microglia modulate blood flow, neurovascular coupling, and hypoperfusion via purinergic actions. Journal of Experimental Medicine, 219(3).

Daftuar, L., Zhu, Y., Jacq, X., & Prives, C. (2013). Ribosomal proteins RPL37, RPS15 and RPS20 regulate the Mdm2-p53-MdmX network. PloS one, 8(7), e68667.

Deczkowska, A., Keren-Shaul, H., Weiner, A., Colonna, M., Schwartz, M., & Amit, I. (2018). Disease-Associated Microglia: A Universal Immune Sensor of Neurodegeneration. Cell, 173(5), 1073–1081. doi:10.1016/j.cell.2018.05.003

Deczkowska, A., Matcovitch-Natan, O., Tsitsou-Kampeli, A., Ben-Hamo, S., Dvir-Szternfeld, R., Spinrad, A., … Smith, L. K. (2017). Mef2C restrains microglial inflammatory response and is lost in brain ageing in an IFN-I-dependent manner. Nature communications, 8(1), 717.

Delhon, L., Mahaut, C., Goudin, N., Gaudas, E., Piquand, K., Le Goff, W., … Le Goff, C. (2019). Impairment of chondrogenesis and microfibrillar network in Adamtsl2 deficiency. The FASEB Journal, 33(2), 2707–2718.

Diaz-Aparicio, I., Paris, I., Sierra-Torre, V., Plaza-Zabala, A., Rodríguez-Iglesias, N., Márquez-Ropero, M., … Alberdi, E. (2020). Microglia actively remodel adult hippocampal neurogenesis through the phagocytosis secretome. Journal of Neuroscience, 40(7), 1453–1482.

Dixon, S. J., Lemberg, K. M., Lamprecht, M. R., Skouta, R., Zaitsev, E. M., Gleason, C. E., … Yang, W. S. (2012). Ferroptosis: an iron-dependent form of nonapoptotic cell death. Cell, 149(5), 1060–1072.

Dobin, A., Davis, C. A., Schlesinger, F., Drenkow, J., Zaleski, C., Jha, S., … Gingeras, T. R. (2013). STAR: ultrafast universal RNA-seq aligner. Bioinformatics, 29(1), 15–21.

Drew, Y., Zenke, F. T., & Curtin, N. J. (2025). DNA damage response inhibitors in cancer therapy: lessons from the past, current status and future implications. Nature Reviews Drug Discovery, 24(1), 19–39.

Eyo, U. B., Peng, J., Swiatkowski, P., Mukherjee, A., Bispo, A., & Wu, L.-J. (2014). Neuronal hyperactivity recruits microglial processes via neuronal NMDA receptors and microglial P2Y12 receptors after status epilepticus. Journal of Neuroscience, 34(32), 10528–10540.

Fagagna, F. d. A. d., Reaper, P. M., Clay-Farrace, L., Fiegler, H., Carr, P., Von Zglinicki, T., … Jackson, S. P. (2003). A DNA damage checkpoint response in telomere-initiated senescence. Nature, 426(6963), 194–198.

François, A., Terro, F., Quellard, N., Fernandez, B., Chassaing, D., Janet, T., … Page, G. (2014). Impairment of autophagy in the central nervous system during lipopolysaccharide-induced inflammatory stress in mice. Molecular brain, 7, 1–16.

Fu, Z., Ganesana, M., Hwang, P., Tan, X., Kinkaid, M. M., Sun, Y. Y., … Kuan, C. Y. (2025). Microglia modulate the cerebrovascular reactivity through ectonucleotidase CD39. Nat Commun, 16(1), 956. doi:10.1038/s41467-025-56093-5

Gandhi, C., Khan, M. M., Lentz, S. R., & Chauhan, A. K. (2012). ADAMTS13 reduces vascular inflammation and the development of early atherosclerosis in mice. *Blood*, The Journal of the American Society of Hematology, 119(10), 2385–2391.

Gao, W., Zhu, J., Westfield, L. A., Tuley, E. A., Anderson, P. J., & Sadler, J. E. (2012). Rearranging exosites in noncatalytic domains can redirect the substrate specificity of ADAMTS proteases. Journal of Biological Chemistry, 287(32), 26944–26952.

Garcia, C., Maurel-Ribes, A., Nauze, M., N’Guyen, D., Martinez, L. O., Payrastre, B., … Pons, V. (2019). Deciphering biased inverse agonism of cangrelor and ticagrelor at P2Y 12 receptor. Cellular and Molecular Life Sciences, 76, 561–576.

Ge, S. X., Jung, D., & Yao, R. (2020). ShinyGO: a graphical gene-set enrichment tool for animals and plants. Bioinformatics, 36(8), 2628–2629.

Geirsdottir, L., David, E., Keren-Shaul, H., Weiner, A., Bohlen, S. C., Neuber, J., … Dutertre, C.-A. (2019). Cross-species single-cell analysis reveals divergence of the primate microglia program. Cell, 179(7), 1609–1622. e1616.

Ginhoux, F., Greter, M., Leboeuf, M., Nandi, S., See, P., Gokhan, S., … Merad, M. (2010). Fate Mapping Analysis Reveals That Adult Microglia Derive from Primitive Macrophages. Science, 330(6005), 841–845. doi:10.1126/science.1194637

Ginhoux, F., & Prinz, M. (2015). Origin of microglia: current concepts and past controversies. Cold Spring Harb Perspect Biol, 7(8), a020537. doi:10.1101/cshperspect.a020537

Gire, V., Roux, P., Wynford-Thomas, D., Brondello, J. M., & Dulic, V. (2004). DNA damage checkpoint kinase Chk2 triggers replicative senescence. The EMBO journal, 23(13), 2554–2563.

Gómez Morillas, A., Besson, V. C., & Lerouet, D. (2021). Microglia and neuroinflammation: what place for P2RY12? International journal of molecular sciences, 22(4), 1636.

Gosselin, D., Skola, D., Coufal, N. G., Holtman, I. R., Schlachetzki, J. C., Sajti, E., … Pasillas, M. P. (2017). An environment-dependent transcriptional network specifies human microglia identity. Science, 356(6344), eaal3222.

Guneykaya, D., Ivanov, A., Hernandez, D. P., Haage, V., Wojtas, B., Meyer, N., … Gielniewski, B. (2018). Transcriptional and translational differences of microglia from male and female brains. Cell reports, 24(10), 2773–2783. e2776.

Hamilton, G., Colbert, J. D., Schuettelkopf, A. W., & Watts, C. (2008). Cystatin F is a cathepsin C-directed protease inhibitor regulated by proteolysis. The EMBO journal, 27(3), 499–508.

Hanisch, U.-K., & Kettenmann, H. (2007). Microglia: active sensor and versatile effector cells in the normal and pathologic brain. Nature Neuroscience, 10(11), 1387–1394.

Hasselmann, J., Coburn, M. A., England, W., Velez, D. X. F., Shabestari, S. K., Tu, C. H., … Claes, C. (2019). Development of a chimeric model to study and manipulate human microglia in vivo. Neuron, 103(6), 1016–1033. e1010.

Haure-Mirande, J.-V., Audrain, M., Ehrlich, M. E., & Gandy, S. (2022). Microglial TYROBP/DAP12 in Alzheimer’s disease: Transduction of physiological and pathological signals across TREM2. Molecular neurodegeneration, 17(1), 55.

Haynes, S. E., Hollopeter, G., Yang, G., Kurpius, D., Dailey, M. E., Gan, W.-B., & Julius, D. (2006). The P2Y12 receptor regulates microglial activation by extracellular nucleotides. Nature neuroscience, 9(12), 1512–1519.

Hickman, S. E., Kingery, N. D., Ohsumi, T. K., Borowsky, M. L., Wang, L.-c., Means, T. K., & El Khoury, J. (2013). The microglial sensome revealed by direct RNA sequencing. Nature Neuroscience, 16(12), 1896–1905.

Holmes, C., Cunningham, C., Zotova, E., Woolford, J., Dean, C., Kerr, S. u., … Perry, V. (2009). Systemic inflammation and disease progression in Alzheimer disease. Neurology, 73(10), 768–774.

Houldsworth, A. (2024). Role of oxidative stress in neurodegenerative disorders: a review of reactive oxygen species and prevention by antioxidants. Brain Communications, 6(1), fcad356.

Iwama, H., Mehanna, S., Imasaka, M., Hashidume, S., Nishiura, H., Yamamura, K.-i., … Ohmuraya, M. (2021). Cathepsin B and D deficiency in the mouse pancreas induces impaired autophagy and chronic pancreatitis. Scientific reports, 11(1), 6596.

Jiang, L., Kon, N., Li, T., Wang, S.-J., Su, T., Hibshoosh, H., … Gu, W. (2015). Ferroptosis as a p53-mediated activity during tumour suppression. Nature, 520(7545), 57–62.

Jung, H., Lee, D., You, H., Lee, M., Kim, H., Cheong, E., & Um, J. W. (2023). LPS induces microglial activation and GABAergic synaptic deficits in the hippocampus accompanied by prolonged cognitive impairment. Scientific reports, 13(1), 6547.

Kenkhuis, B., Somarakis, A., Kleindouwel, L. R. T., van Roon-Mom, W. M. C., Hollt, T., & van der Weerd, L. (2022). Co-expression patterns of microglia markers Iba1, TMEM119 and P2RY12 in Alzheimer’s disease. Neurobiol Dis, 167, 105684. doi:10.1016/j.nbd.2022.105684

Keren-Shaul, H., Spinrad, A., Weiner, A., Matcovitch-Natan, O., Dvir-Szternfeld, R., Ulland, T., … Toth, B. (2017). A unique microglia type associated with restricting development of Alzheimer’s disease. 1276–1290 e1217 Cell 169. In.

Keren-Shaul, H., Spinrad, A., Weiner, A., Matcovitch-Natan, O., Dvir-Szternfeld, R., Ulland, T. K., … Amit, I. (2017). A Unique Microglia Type Associated with Restricting Development of Alzheimer’s Disease. Cell, 169(7), 1276–1290 e1217. doi:10.1016/j.cell.2017.05.018

Kettenmann, H., Hanisch, U.-K., Noda, M., & Verkhratsky, A. (2011). Physiology of microglia. Physiological reviews, 91(2), 461–553.

Kim, S. E., Zhang, L., Ma, K., Riegman, M., Chen, F., Ingold, I., … Jiang, X. (2016). Ultrasmall nanoparticles induce ferroptosis in nutrient-deprived cancer cells and suppress tumour growth. Nature nanotechnology, 11(11), 977–985.

Klaassen, C. D., Liu, J., & Choudhuri, S. (1999). Metallothionein: an intracellular protein to protect against cadmium toxicity. Annual review of pharmacology and toxicology, 39(1), 267–294.

Kopec, K. K., & Carroll, R. T. (2000). Phagocytosis is regulated by nitric oxide in murine microglia. Nitric Oxide, 4(2), 103–111.

Kruse, J.-P., & Gu, W. (2009). Modes of p53 regulation. Cell, 137(4), 609–622.

Langfelder, P., & Horvath, S. (2008). WGCNA: an R package for weighted correlation network analysis. BMC Bioinformatics, 9(1), 559. doi:10.1186/1471-2105-9-559

Laroux, F. S., Romero, X., Wetzler, L., Engel, P., & Terhorst, C. (2005). Cutting edge: MyD88 controls phagocyte NADPH oxidase function and killing of gram-negative bacteria. The Journal of Immunology, 175(9), 5596–5600.

Lavin, Y., Winter, D., Blecher-Gonen, R., David, E., Keren-Shaul, H., Merad, M., … Amit, I. (2014). Tissue-resident macrophage enhancer landscapes are shaped by the local microenvironment. Cell, 159(6), 1312–1326.

Lazo, J. S., & Pitt, B. R. (1995). Metallothioneins and cell death by anticancer drugs. Annual review of pharmacology and toxicology, 35, 635–653.

Le Goff, C., Morice-Picard, F., Dagoneau, N., Wang, L. W., Perrot, C., Crow, Y. J., … Krakow, D. (2008). ADAMTSL2 mutations in geleophysic dysplasia demonstrate a role for ADAMTS-like proteins in TGF-β bioavailability regulation. Nature genetics, 40(9), 1119–1123.

Lei, S., & Liu, Y. (2024). Identifying microglia-derived NFKBIA as a potential contributor to the pathogenesis of Alzheimer’s disease and age-related macular degeneration. Journal of Alzheimer’s Disease, 13872877251326267.

Li, W., Garringer, H. J., Goodwin, C. B., Richine, B., Acton, A., VanDuyn, N., … Peacock, M. (2015). Systemic and cerebral iron homeostasis in ferritin knock-out mice. PloS one, 10(1), e0117435.

Li, X., Wu, L., Sun, L., Liu, H., Qiao, X., Mi, N., … Quan, P. (2024). Ferroptosis-Related Gene Signatures in Epilepsy: Diagnostic and Immune Insights. Molecular Neurobiology, 1–14.

Liddell, J. R., Hilton, J. B., Kysenius, K., Billings, J. L., Nikseresht, S., McInnes, L. E., … Belaidi, A. A. (2024). Microglial ferroptotic stress causes non-cell autonomous neuronal death. Molecular neurodegeneration, 19(1), 14.

Lin, Y., Yang, Q., Lin, X., Liu, X., Qian, Y., Xu, D., … Hu, W. (2024). Extracellular matrix disorganization caused by ADAMTS16 deficiency leads to bicuspid aortic valve with raphe formation. Circulation, 149(8), 605–626.

Linkermann, A., Skouta, R., Himmerkus, N., Mulay, S. R., Dewitz, C., De Zen, F., … Welz, P.-S. (2014). Synchronized renal tubular cell death involves ferroptosis. Proceedings of the National Academy of Sciences, 111(47), 16836–16841.

Liu, Q., Zhang, Y., Liu, S., Liu, Y., Yang, X., Liu, G., … Ma, J. (2019). Cathepsin C promotes microglia M1 polarization and aggravates neuroinflammation via activation of Ca 2+-dependent PKC/p38MAPK/NF-κB pathway. Journal of neuroinflammation, 16, 1–18.

Liu, S., Gao, X., & Zhou, S. (2022). New target for prevention and treatment of neuroinflammation: microglia iron accumulation and ferroptosis. ASN neuro, 14, 17590914221133236.

Liu, Y., & Gu, W. (2022). p53 in ferroptosis regulation: the new weapon for the old guardian. Cell Death & Differentiation, 29(5), 895–910.

Liu, Y., Wu, J., Najem, H., Lin, Y., Pang, L., Khan, F., … Chen, P. (2024). Dual targeting macrophages and microglia is a therapeutic vulnerability in models of PTEN-deficient glioblastoma. The Journal of Clinical Investigation, 134(22).

Lively, S., & Schlichter, L. C. (2018). Microglia responses to pro-inflammatory stimuli (LPS, IFNγ+ TNFα) and reprogramming by resolving cytokines (IL-4, IL-10). Front Cell Neurosci, 12, 215.

Lopes, K. O., Sparks, D. L., & Streit, W. J. (2008). Microglial dystrophy in the aged and Alzheimer’s disease brain is associated with ferritin immunoreactivity. Glia, 56(10), 1048–1060.

Love, M. I., Huber, W., & Anders, S. (2014). Moderated estimation of fold change and dispersion for RNA-seq data with DESeq2. Genome biology, 15, 1–21.

Lupica-Tondo, G. L., Arner, E. N., Mogilenko, D. A., & Voss, K. (2024). Immunometabolism of ferroptosis in the tumor microenvironment. Frontiers in Oncology, 14, 1441338.

Marsh, S. E., Walker, A. J., Kamath, T., Dissing-Olesen, L., Hammond, T. R., de Soysa, T. Y., … Nadaf, N. (2022). Dissection of artifactual and confounding glial signatures by single-cell sequencing of mouse and human brain. Nature Neuroscience, 25(3), 306–316.

Martin, M. (2011). Cutadapt removes adapter sequences from high-throughput sequencing reads. *EMBnet*. journal, 17(1), 10–12.

Masuda, T., Sankowski, R., Staszewski, O., Böttcher, C., Amann, L., Sagar, N., … van Loo, G. (2019). Spatial and temporal heterogeneity of mouse and human microglia at single-cell resolution. Nature, 566(7744), 388–392.

McCarthy, R. C., Sosa, J. C., Gardeck, A. M., Baez, A. S., Lee, C.-H., & Wessling-Resnick, M. (2018). Inflammation-induced iron transport and metabolism by brain microglia. Journal of Biological Chemistry, 293(20), 7853–7863.

Mildner, A., Huang, H., Radke, J., Stenzel, W., & Priller, J. (2017). P2Y12 receptor is expressed on human microglia under physiological conditions throughout development and is sensitive to neuroinflammatory diseases. Glia, 65(2), 375–387.

Miller, S. C., MacDonald, C. C., Kellogg, M. K., Karamysheva, Z. N., & Karamyshev, A. L. (2023). Specialized ribosomes in health and disease. International journal of molecular sciences, 24(7), 6334.

Monsonego, A., Imitola, J., Zota, V., Oida, T., & Weiner, H. L. (2003). Microglia-mediated nitric oxide cytotoxicity of T cells following amyloid β-peptide presentation to Th1 cells. The Journal of Immunology, 171(5), 2216–2224.

Moore, C. S., Ase, A. R., Kinsara, A., Rao, V. T., Michell-Robinson, M., Leong, S. Y., … Bar-Or, A. (2015). P2Y12 expression and function in alternatively activated human microglia. Neuroimmunology & Neuroinflammation, 2(2), e80.

Nakanishi, H., Zhang, J., Koike, M., Nishioku, T., Okamoto, Y., Kominami, E., … Saftig, P. (2001). Involvement of nitric oxide released from microglia–macrophages in pathological changes of cathepsin D-deficient mice. Journal of Neuroscience, 21(19), 7526–7533.

Nie, H., Ju, H., Fan, J., Shi, X., Cheng, Y., Cang, X., … Yi, W. (2020). O-GlcNAcylation of PGK1 coordinates glycolysis and TCA cycle to promote tumor growth. Nature communications, 11(1), 36.

Nishioku, T., Hashimoto, K., Yamashita, K., Liou, S.-Y., Kagamiishi, Y., Maegawa, H., … Saftig, P. (2002). Involvement of cathepsin E in exogenous antigen processing in primary cultured murine microglia. Journal of Biological Chemistry, 277(7), 4816–4822.

Obermeier, B., Daneman, R., & Ransohoff, R. M. (2013). Development, maintenance and disruption of the blood-brain barrier. Nature medicine, 19(12), 1584–1596.

Palmer, C. R., Liu, C. S., Romanow, W. J., Lee, M.-H., & Chun, J. (2021). Altered cell and RNA isoform diversity in aging Down syndrome brains. Proceedings of the National Academy of Sciences, 118(47), e2114326118.

Paolicelli, R. C., Bolasco, G., Pagani, F., Maggi, L., Scianni, M., Panzanelli, P., … Dumas, L. (2011). Synaptic pruning by microglia is necessary for normal brain development. Science, 333(6048), 1456–1458.

Paolicelli, R. C., Sierra, A., Stevens, B., Tremblay, M.-E., Aguzzi, A., Ajami, B., … Bennett, M. (2022). Microglia states and nomenclature: A field at its crossroads. Neuron, 110(21), 3458–3483.

Peng, J., Liu, Y., Umpierre, A. D., Xie, M., Tian, D.-S., Richardson, J. R., & Wu, L.-J. (2019). Microglial P2Y12 receptor regulates ventral hippocampal CA1 neuronal excitability and innate fear in mice. Molecular brain, 12, 1–10.

Peng, L.-Y., Wang, J., Tao, M., You, C.-P., Ye, L., Xiao, J., … Liu, S.-J. (2014). Analysis of Mitochondrial Respiratory-Related Genes Reveals Nuclear and Mitochondrial Genome Cooperation in Allotetraploid Hybrid. Current Molecular Medicine, 14(10), 1314–1321.

Perry, V. H., & Holmes, C. (2014). Microglial priming in neurodegenerative disease. Nature Reviews Neurology, 10(4), 217–224.

Perry, V. H., Newman, T. A., & Cunningham, C. (2003). The impact of systemic infection on the progression of neurodegenerative disease. Nature Reviews Neuroscience, 4(2), 103–112.

Perry, V. H., Nicoll, J. A., & Holmes, C. (2010). Microglia in neurodegenerative disease. Nature Reviews Neurology, 6(4), 193–201.

Rangaraju, S., Dammer, E. B., Raza, S. A., Rathakrishnan, P., Xiao, H., Gao, T., … Seyfried, N. T. (2018). Identification and therapeutic modulation of a pro-inflammatory subset of disease-associated-microglia in Alzheimer’s disease. Molecular neurodegeneration, 13, 1–25.

Ransohoff, R. M., & Perry, V. H. (2009). Microglial physiology: unique stimuli, specialized responses. Annual review of immunology, 27(1), 119–145.

Reddy, K. R. K., Dasari, C., Duscharla, D., Supriya, B., Ram, N. S., Surekha, M., … Ummanni, R. (2018). Dimethylarginine dimethylaminohydrolase-1 (DDAH1) is frequently upregulated in prostate cancer, and its overexpression conveys tumor growth and angiogenesis by metabolizing asymmetric dimethylarginine (ADMA). Angiogenesis, 21, 79–94.

Riegman, M., Sagie, L., Galed, C., Levin, T., Steinberg, N., Dixon, S. J., … Zaritsky, A. (2020). Ferroptosis occurs through an osmotic mechanism and propagates independently of cell rupture. Nature cell biology, 22(9), 1042–1048.

Sabogal-Guáqueta, A. M., Marmolejo-Garza, A., Trombetta-Lima, M., Oun, A., Hunneman, J., Chen, T., … Wolters, J. C. (2023). Species-specific metabolic reprogramming in human and mouse microglia during inflammatory pathway induction. Nature communications, 14(1), 6454.

Salter, M. W., & Stevens, B. (2017). Microglia emerge as central players in brain disease. Nat Med, 23(9), 1018–1027. doi:10.1038/nm.4397

Schioppa, T., Sozio, F., Barbazza, I., Scutera, S., Bosisio, D., Sozzani, S., & Del Prete, A. (2020). Molecular basis for CCRL2 regulation of leukocyte migration. Frontiers in Cell and Developmental Biology, 8, 615031.

Seibt, T. M., Proneth, B., & Conrad, M. (2019). Role of GPX4 in ferroptosis and its pharmacological implication. Free radical biology and medicine, 133, 144–152.

Sfera, A., Gradini, R., Cummings, M., Diaz, E., Price, A. I., & Osorio, C. (2018). Rusty microglia: trainers of innate immunity in Alzheimer’s disease. Frontiers in neurology, 9, 1062.

Shannon, P., Markiel, A., Ozier, O., Baliga, N. S., Wang, J. T., Ramage, D., … Ideker, T. (2003). Cytoscape: a software environment for integrated models of biomolecular interaction networks. Genome research, 13(11), 2498–2504.

Sharma, L. K., Lu, J., & Bai, Y. (2009). Mitochondrial respiratory complex I: structure, function and implication in human diseases. Current medicinal chemistry, 16(10), 1266–1277.

Shin, J. A., Kim, Y. A., Kim, H. W., Kim, H.-S., Lee, K.-E., Kang, J. L., & Park, E.-M. (2018). Iron released from reactive microglia by noggin improves myelin repair in the ischemic brain. Neuropharmacology, 133, 202–215.

Sierra, A., Encinas, J. M., Deudero, J. J., Chancey, J. H., Enikolopov, G., Overstreet-Wadiche, L. S., … Maletic-Savatic, M. (2010). Microglia shape adult hippocampal neurogenesis through apoptosis-coupled phagocytosis. Cell stem cell, 7(4), 483–495.

Sierra, A., Miron, V. E., Paolicelli, R. C., & Ransohoff, R. M. (2024). Microglia in Health and Diseases: Integrative Hubs of the Central Nervous System (CNS). *Cold Spring Harbor Perspectives in Biology*, a041366.

Škandík, M., Friess, L., Vázquez-Cabrera, G., Keane, L., Grabert, K., Cruz De los Santos, M., … Joseph, B. (2025). Age-associated microglial transcriptome leads to diminished immunogenicity and dysregulation of MCT4 and P2RY12/P2RY13 related functions. Cell Death Discovery, 11(1), 16.

Smith, L. K., He, Y., Park, J.-S., Bieri, G., Snethlage, C. E., Lin, K., … Udeochu, J. (2015). β2-microglobulin is a systemic pro-aging factor that impairs cognitive function and neurogenesis. Nature medicine, 21(8), 932–937.

Socodato, R., & Relvas, J. o. B. (2025). Neuroinflammation revisited through the microglial lens. Neural Regeneration Research, 20(7), 1989–1990.

Stanton, H., Melrose, J., Little, C. B., & Fosang, A. J. (2011). Proteoglycan degradation by the ADAMTS family of proteinases. Biochimica et Biophysica Acta (BBA)-Molecular Basis of Disease, 1812(12), 1616–1629.

Stockwell, B. R., Angeli, J. P. F., Bayir, H., Bush, A. I., Conrad, M., Dixon, S. J., … Kagan, V. E. (2017). Ferroptosis: a regulated cell death nexus linking metabolism, redox biology, and disease. Cell, 171(2), 273–285.

Strohm, A. O., Oldfield, S., Hernady, E., Johnston, C. J., Marples, B., O’Banion, M. K., & Majewska, A. K. (2025). Biological sex, microglial signaling pathways, and radiation exposure shape cortical proteomic profiles and behavior in mice. *Brain, Behavior*, & Immunity-Health, 43, 100911.

Tremblay, M.-È., Stevens, B., Sierra, A., Wake, H., Bessis, A., & Nimmerjahn, A. (2011). The role of microglia in the healthy brain. Journal of Neuroscience, 31(45), 16064–16069.

Umpierre, A. D., & Wu, L. J. (2021). How microglia sense and regulate neuronal activity. Glia, 69(7), 1637–1653.

Urrutia, P., Aguirre, P., Esparza, A., Tapia, V., Mena, N. P., Arredondo, M., … Núñez, M. T. (2013). Inflammation alters the expression of DMT 1, FPN 1 and hepcidin, and it causes iron accumulation in central nervous system cells. Journal of neurochemistry, 126(4), 541–549.

van Wageningen, T. A., Vlaar, E., Kooij, G., Jongenelen, C. A. M., Geurts, J. J. G., & van Dam, A. M. (2019). Regulation of microglial TMEM119 and P2RY12 immunoreactivity in multiple sclerosis white and grey matter lesions is dependent on their inflammatory environment. Acta Neuropathol Commun, 7(1), 206. doi:10.1186/s40478-019-0850-z

Vezzani, A., Fujinami, R. S., White, H. S., Preux, P.-M., Blümcke, I., Sander, J. W., & Löscher, W. (2016). Infections, inflammation and epilepsy. Acta neuropathologica, 131(2), 211–234.

Villa, A., Gelosa, P., Castiglioni, L., Cimino, M., Rizzi, N., Pepe, G., … Vegeto, E. (2018). Sex-specific features of microglia from adult mice. Cell reports, 23(12), 3501–3511.

Walker, D. G., Tang, T. M., Mendsaikhan, A., Tooyama, I., Serrano, G. E., Sue, L. I., … Lue, L.-F. (2020). Patterns of expression of purinergic receptor P2RY12, a putative marker for non-activated microglia, in aged and Alzheimer’s disease brains. International journal of molecular sciences, 21(2), 678.

Wang, C., Yue, H., Hu, Z., Shen, Y., Ma, J., Li, J., … Shi, P. (2020). Microglia mediate forgetting via complement-dependent synaptic elimination. Science, 367(6478), 688–694.

Wang, Q., Shi, N.-R., Lv, P., Liu, J., Zhang, J.-Z., Deng, B.-L., … Chen, X. (2023). P2Y12 receptor gene polymorphisms are associated with epilepsy. Purinergic Signalling, 19(1), 155–162.

Yang, Y., Wang, Y., Guo, L., Gao, W., Tang, T.-L., & Yan, M. (2022). Interaction between macrophages and ferroptosis. Cell death & disease, 13(4), 355.

Ye, C., Huang, X., Tong, Y., Chen, Y., Zhao, X., Xie, W., … Mo, Y. (2025). Overexpression of ALKBH5 alleviates LPS induced neuroinflammation via increasing NFKBIA. International immunopharmacology, 144, 113598.

Yu, G., Wang, L.-G., Han, Y., & He, Q.-Y. (2012). clusterProfiler: an R package for comparing biological themes among gene clusters. Omics: a journal of integrative biology, 16(5), 284–287.

Yu, X., & Long, Y. C. (2016). Crosstalk between cystine and glutathione is critical for the regulation of amino acid signaling pathways and ferroptosis. Scientific reports, 6(1), 30033.

Zhang, M., Zhou, Y., Jiang, Y., Lu, Z., Xiao, X., Ning, J., … Can, D. (2021). Profiling of sexually dimorphic genes in neural cells to identify Eif2s3y, whose overexpression causes autism-like behaviors in male mice. Frontiers in Cell and Developmental Biology, 9, 669798.

Zhang, N., Yu, X., Xie, J., & Xu, H. (2021). New insights into the role of ferritin in iron homeostasis and neurodegenerative diseases. Molecular Neurobiology, 58(6), 2812–2823.

Zhang, T., Ding, C., Chen, H., Zhao, J., Chen, Z., Chen, B., … Xu, H. (2022). m6A mRNA modification maintains colonic epithelial cell homeostasis via NF-κB–mediated antiapoptotic pathway. Science advances, 8(12), eabl5723.

Zhao, G., Arosio, P., & Chasteen, N. D. (2006). Iron (II) and hydrogen peroxide detoxification by human H-chain ferritin. An EPR spin-trapping study. Biochemistry, 45(10), 3429–3436.

Zhao, H., Wang, J., Li, Z., Wang, S., Yu, G., & Wang, L. (2023). Identification ferroptosis-related hub genes and diagnostic model in Alzheimer’s disease. Frontiers in Molecular Neuroscience, 16, 1280639.

Zhao, K., Sun, Y., Zhong, S., & Luo, J.-L. (2024). The multifaceted roles of cathepsins in immune and inflammatory responses: implications for cancer therapy, autoimmune diseases, and infectious diseases. Biomarker Research, 12(1), 165.

Zrzavy, T., Hametner, S., Wimmer, I., Butovsky, O., Weiner, H. L., & Lassmann, H. (2017). Loss of ‘homeostatic’microglia and patterns of their activation in active multiple sclerosis. Brain, 140(7), 1900–1913.

